# The Effects of Locus Coeruleus Optogenetic Stimulation on Global Spatiotemporal Patterns in Rats

**DOI:** 10.1101/2024.05.23.595327

**Authors:** Nmachi Anumba, Michael A. Kelberman, Wenju Pan, Alexia Marriott, Xiaodi Zhang, Nan Xu, David Weinshenker, Shella Keilholz

## Abstract

Whole-brain intrinsic activity as detected by resting-state fMRI can be summarized by three primary spatiotemporal patterns. These patterns have been shown to change with different brain states, especially arousal. The noradrenergic locus coeruleus (LC) is a key node in arousal circuits and has extensive projections throughout the brain, giving it neuromodulatory influence over the coordinated activity of structurally separated regions. In this study, we used optogenetic-fMRI in rats to investigate the impact of LC stimulation on the global signal and three primary spatiotemporal patterns. We report small, spatially specific changes in global signal distribution as a result of tonic LC stimulation, as well as regional changes in spatiotemporal patterns of activity at 5 Hz tonic and 15 Hz phasic stimulation. We also found that LC stimulation had little to no effect on the spatiotemporal patterns detected by complex principal component analysis. These results show that the effects of LC activity on the BOLD signal in rats may be small and regionally concentrated, as opposed to widespread and globally acting.

## 1. INTRODUCTION

Resting-state functional MRI (rs-fMRI) is increasingly used to non-invasively study dynamic brain activity and the intrinsic forms of large-scale communication across the brain. Of particular interest are recent studies that have shown a majority of spatially structured signals captured using rs-fMRI can be summarized or represented by three primary spatiotemporal patterns, also known as quasiperiodic patterns or QPPs (Bolt et al., 2022; Majeed et al., 2011; Yousefi et al., 2018; Yousefi & Keilholz, 2021). The most prominent of these patterns, QPP1, captures the spatiotemporal evolution of the global blood oxygenated level dependent (BOLD) signal. The global signal, typically measured to be the average of all brain activity over the course of a scan, is often used as a nuisance regressor to reduce the effects of widespread noise in rs-fMRI data (Birn et al., 2006; Parkes et al., 2018; Satterthwaite et al., 2012; Yan et al., 2013); although the existence of resting-state networks and large-scale patterns of activity have caused many to question the removal of the global signal from fMRI datasets (Ciric et al., 2017; Fox et al., 2009; T. T. Liu et al., 2017; Murphy & Fox, 2017; Saad et al., 2012). The second strongest spatiotemporal pattern, QPP2, is comprised of semi-regular waves of alternating activity between different structures and networks that have been observed across species. In humans, QPP2 captures a cyclical pattern of anticorrelation between the default mode network (DMN) and task positive network (TPN), along with propagation across the cortex during the transition between activation of the two networks (Majeed et al., 2011). In rodents, it manifests as anticorrelation between the rodent DMN and lateral cortical network, the latter of which is believed to exhibit similar activity to the human TPN (Belloy et al., 2018). The anticorrelated QPP also contributes substantially to functional connectivity across the whole brain (Abbas, Belloy, et al., 2019; Belloy et al., 2018; Yousefi et al., 2018). Of note that most prior studies employed global signal regression, making QPP2 the primary spatiotemporal pattern that was observed.

Despite most often being measured during resting state, both of these spatiotemporal patterns exhibit notable changes during arousal and in correspondence with vigilance-related measures. Specifically, the global signal (also captured by QPP1) is inversely correlated with various measures of vigilance and arousal across different species. In humans, an EEG measurement of vigilance was found to be inversely correlated with the amplitude of the global signal (Wong et al., 2013, 2016). A negative correlation to the global signal was also found with a local field potential measure of arousal in non-human primates (Chang et al., 2016). A separate study in mice used pupil diameter as a proxy for arousal and showed that changes in pupil diameter were anticorrelated to changes in global hemodynamics (Pisauro et al., 2016). Additionally, the introduction of caffeine as a measure of vigilance has been shown to increase anti-correlation between the DMN and the TPN (Wong et al., 2012), and a wave-like propagation of phase shifts was reported to occur as a result of infra-slow arousal-related activity (Raut et al., 2021). These relationships suggest that the global signal and QPP1 typically increase in strength as arousal diminishes.

QPP2 also has ties to attention and arousal. Early work showed that time-varying anticorrelation between the DMN and TPN was related to intra– and inter-individual variability in reaction time on a psychomotor vigilance task (Thompson et al., 2013). Further analysis showed that these fluctuations in reaction time were related to the phase of QPP2 at task onset (Abbas et al., 2016). The contribution of QPP2 to rs-fMRI is reduced in patients with ADHD compared to healthy controls (Abbas, Bassil, et al., 2019). Taken together, these findings suggest that the strength of QPP2 should increase with increasing arousal. Recent analysis of data from the Human Connectome Project indicates that QPP1 increases at the expense of QPP2 over the course of a scan, consistent with growing drowsiness and prior work that examined QPP1 and QPP2 separately (Meyer-Baese et al., 2024).

The brainstem noradrenergic locus coeruleus (LC) is a key node in the brain’s arousal system and regulates sleep, stress responses, mood, and cognition (Aston-Jones & Bloom, 1981; Aston-Jones & Cohen, 2005; Benarroch, 2018; Poe et al., 2020; Rajkowski et al., 1994). In recent years, the LC-norepinephrine (LC-NE) system has been recognized as an important contributor to the generation of spatiotemporal dynamic signals. Of interest to fMRI in particular, it has also been shown that LC-NE vascular innervation helps to optimize neurovascular coupling, leading to a potential increase in sensitivity of the BOLD signal (Bekar et al., 2012). As the primary source of central NE and potential for brain-wide release, the vast projections of the LC have potential to influence whole-brain levels of activity. In fact, evidence shows that the activity of brainstem nuclei may play a significant role in large-scale spatiotemporal signals (van den Brink et al., 2019). Turchi et al., showed that inhibition of the basal forebrain decreased the amplitude of global signal fluctuations (Turchi et al., 2018). However, these effects were more pronounced during less alert behavioral states, indicating a connection of this effect to arousal and potentially the activity of other brainstem nuclei. The origin of QPP2 has been speculated to arise from the activity of neuromodulatory brainstem nuclei (Abbas, Belloy, et al., 2019), and preliminary work has shown that extreme modulation of NE levels (either up or down) dramatically reduces the detection of this QPP (Abbas et al., 2018). The LC-NE system has also been hypothesized to coordinate and integrate activity across structurally segregated regions (Shine, 2019). LC activation has further been shown to affect large-scale measures of activity by strengthening brain-wide connectivity and even modulating nodes of the rodent DMN (Oyarzabal et al., 2021; Zerbi et al., 2019).

In this study, we aimed to further investigate the contribution of the LC-NE system to the composition of the global signal, QPP1 and QPP2 through the utilization of optogenetic-fMRI. Optogenetics is optimal for this purpose because, beyond general cell-type specific stimulation, it has the capacity to employ specific stimulation rates, patterns, and timing. This is critical for the LC because it demonstrates contrasting firing rates and patterns that are associated with distinct arousal and cognitive states. LC neurons have low tonic pacemaker activity (∼1-2 Hz) during general wakefulness, high tonic activity (∼5-10 Hz) in response to stress, and phasic activity (∼15 Hz bursts followed by inhibition) during focused attention and goal-directed behavior (Aston-Jones & Bloom, 1981; Aston-Jones & Cohen, 2005; Berridge & Waterhouse, 2003; Carter et al., 2010; McCall et al., 2015; Prokopiou et al., 2022; Valentino & Foote, 1988). Thus, we used optogenetic-fMRI in rats to investigate distinct characteristics of the global signal, QPP1 and QPP2 during low tonic (2 Hz), high tonic (5 Hz), and phasic (15 Hz bursts) stimulation of the LC that mimic natural patterns of LC activity. We report small, spatially-specific changes in global signal distribution under tonic LC stimulation, as well as regional changes in QPP2 involvement during 5 Hz tonic and 15 Hz phasic stimulation. We also show little to no effect of LC stimulation on these spatiotemporal patterns as detected by complex principal component analysis (CPCA), an analysis that simultaneously detects the global signal and the anticorrelated QPP from BOLD signals (Bolt et al., 2022). These results indicate that the neuromodulatory effects as a result of LC optogenetic stimulation on large-scale spatiotemporal patterns are relatively small in magnitude and concentrated in particular brain regions.

## 2. MATERIALS AND METHODS

An overview of the methods involved in the optogenetic-fMRI experimentation from initial surgery to data analysis can be seen in **Figure 1**.

**Figure 1:**
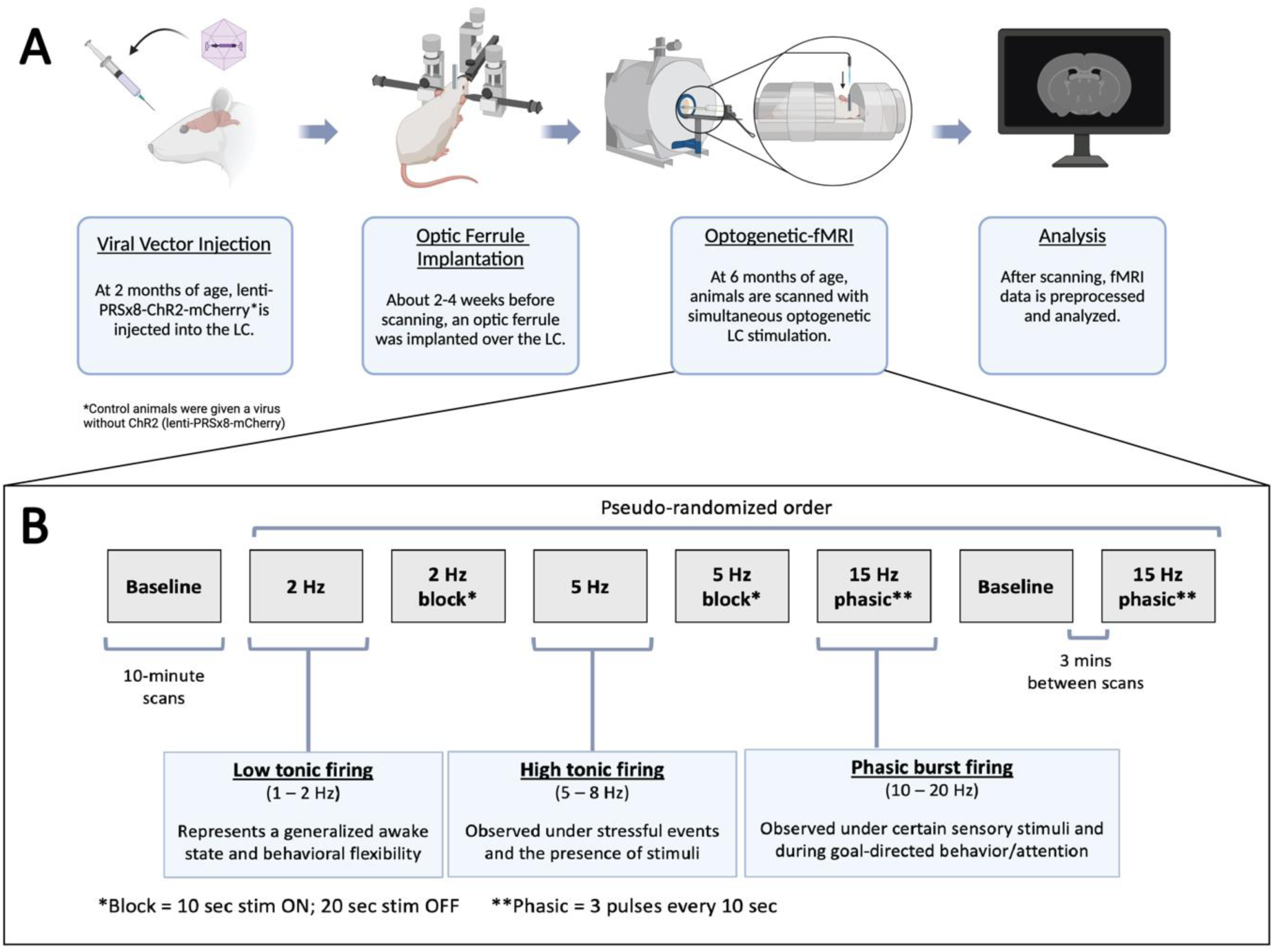
Optogenetic-fMRI experimental methods. (A) Experimental timeline. 2-month-old rats received bilateral injections of a lentivirus containing the synthetic noradrenergic promoter PRSx8, the mCherry fluorescent protein, and channelrhodopsin-2 (ChR2) if they were not control animals. 2-4 weeks before scanning took place, animals were implanted unilaterally with an optic ferrule over the LC. At 6 months of age, animals underwent an fMRI scanning session with varying levels of simultaneous optogenetic LC stimulation. After scanning, all fMRI data was preprocessed and analyzed. (B) Scanning session timeline. All scanning sessions consisted of eight 10-minute fMRI scans with three minutes between scans. The LC was stimulated using 2 Hz, 5 Hz, and 15 Hz phasic bursts to replicate the common modes of firing displayed by the LC. Each session started with a baseline scan (no stimulation) that was followed by the remaining seven scans in a pseudo-randomized order. The block stimulation scans were not used in this study. Created in part with BioRender.com.

### 2.1 Animal Preparation

Two cohorts of 6-month-old male and female wildtype Fischer rats were used in this study. All rats were bred in-house, kept on a 12 hr light/dark cycle (lights on at 7:00 am), and provided access to food and water *ad libitum* throughout the experimental period. Rats were group-housed in groups of 2-3 animals up until implantation of the optic ferrule, after which they were singly housed until they underwent scanning in the MRI machine (approximately 2-4 weeks). Experimental groups were comprised of animals of both sexes. For a breakdown of the sex makeup of each group refer to (**Supplementary Table 1**). For information regarding number of subjects and scans, see **Table 1**. All protocols used in this study were approved by the Institutional Animal Care and Use Committee (IACUC) at Emory University and all procedures were performed in accordance with the IACUC protocols. All surgeries and experiments were carried out under isoflurane anesthesia and all efforts were made to minimize the stress and suffering of the animals.

**Table 1:**
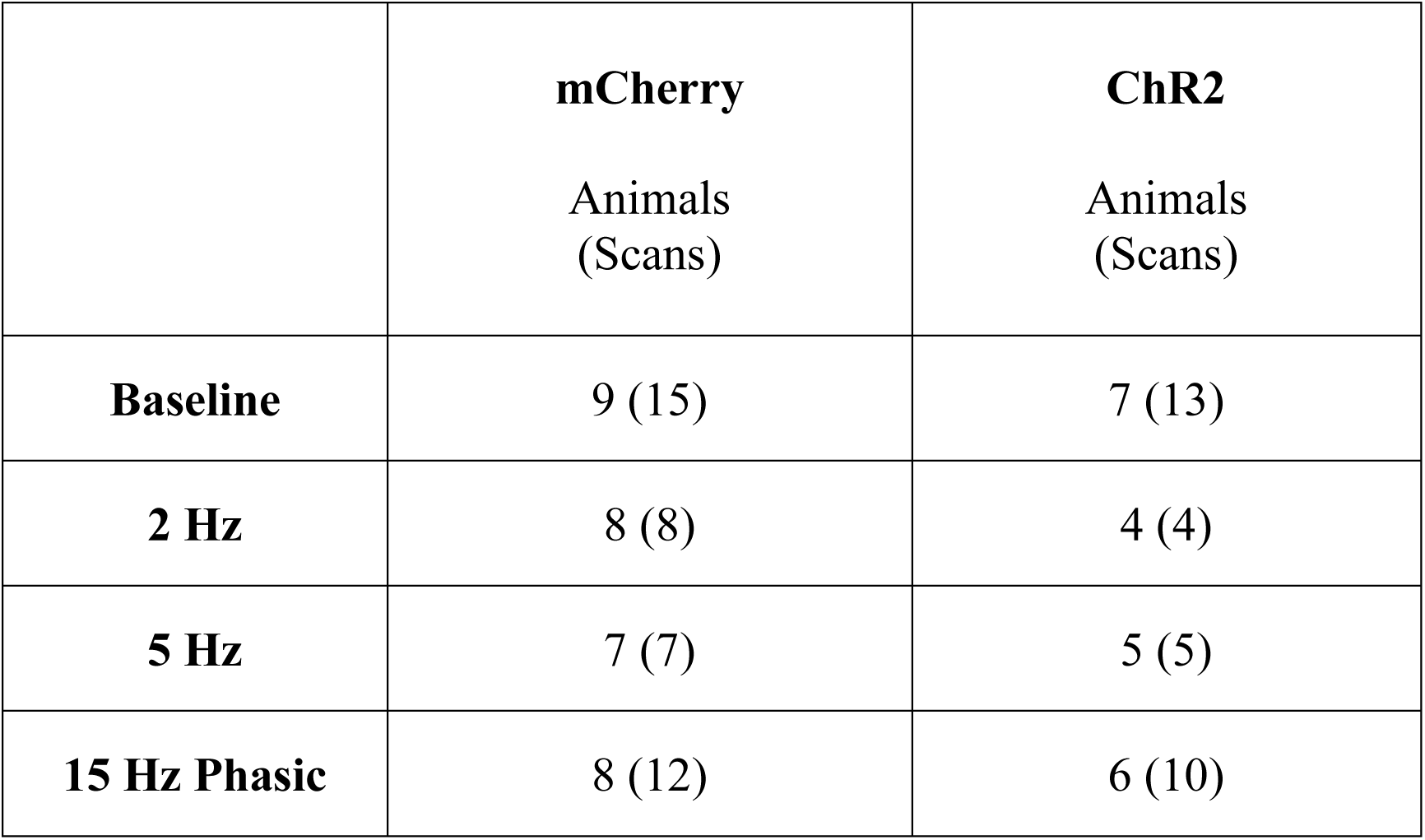
Animal group sizes.

|  | **mCherry**<br>Animals<br>(Scans) | **ChR2**<br>Animals<br>(Scans) |
| --- | --- | --- |
| **Baseline** | 9 (15) | 7 (13) |
| **2 Hz** | 8 (8) | 4 (4) |
| **5 Hz** | 7 (7) | 5 (5) |
| **15 Hz Phasic** | 8 (12) | 6 (10) |

### 2.2 Surgery and Pupillometry

Stereotaxic surgery was performed on all rats at approximately 2-months of age. Rats were initially put under anesthesia with 5% isoflurane. Isoflurane was then lowered to 2% for the remainder of the surgery and rats were given ketoprofen (5 mg/kg, s.c.) directly following incision. To avoid contact with the sagittal sinus a 15-degree head tilt was utilized and bilateral injections (1.3 µl/hemisphere; AP: –3.8 mm from lambda, ML: +/− 1.2 mm, DV: 7.0 mm from skull surface) of a lentivirus under control of the synthetic noradrenergic promoter, PRSx8, was used to express either ChR2-mCherry or mCherry selectively in the LC (Abbott et al., 2009; Hwang et al., 2001; Vazey & Aston-Jones, 2014). After the infusion, the needle was left for 5 minutes after which it was moved 1 mm dorsally and 2 minutes were allotted to permit appropriate viral diffusion at the injection site.

Approximately 2–4 weeks before undergoing MRI scanning rats were implanted with a custom fiber optic ferrule (FG200UCC, 200 um, 0.22 NA, 7.5 mm length; Thorlabs) above the LC to enable delivery of optogenetic stimulation. The hemisphere in which the LC was targeted was randomized for each rat so that stimulation of both the left and right LC were incorporated into group analyses. First, the skull was implanted with three MRI-compatible screws to hold the ferrule in place. The aforementioned surgical protocol was then performed with an optic ferrule being implanted dorsal to the LC (AP: –3.8 mm from lambda, ML: +/−1.2 mm, DV: 6.5-6.8 mm from skull surface).

Stimulation of the LC results in pupil dilation (Y. Liu et al., 2017; Zerbi et al., 2019), providing a useful measure to verify successful viral expression and optic ferrule targeting. We performed this verification via pupillometry following previously published protocols (Privitera et al., 2020). Our setup involved the use of a Raspberry Pi NoIR Camera Module V2 night vision camera, an infrared light source, and a Raspberry Pi 4 Model B (CanaKit). During optic ferrule implantation, 90 seconds of pupil recordings were taken of the eye ipsilateral to the hemisphere of the stimulated LC. The recordings were comprised of 30 seconds of baseline (no stimulation) followed by 30 seconds of LC stimulation, followed by 30 seconds of post-stimulation baseline footage (**Figure 2B**). Pupil dilation was quantified offline using a previously established pipeline (Privitera et al., 2020). In the event that ChR2-mCherry did not display pupil dilation in response to LC stimulation, the other hemisphere was implanted and checked for pupil dilation. If there was still no pupil dilation, animals were re-categorized to the control group. During offline analysis, there were two animals where pupil recording lengths differed. These videos were manually checked by an experimenter blinded to experimental condition for the presence or lack of pupil dilation in response to LC stimulation. Dental cement was used to secure the implant to the skull and animals were allowed to recover for approximately 2-4 weeks.

**Figure 2:**
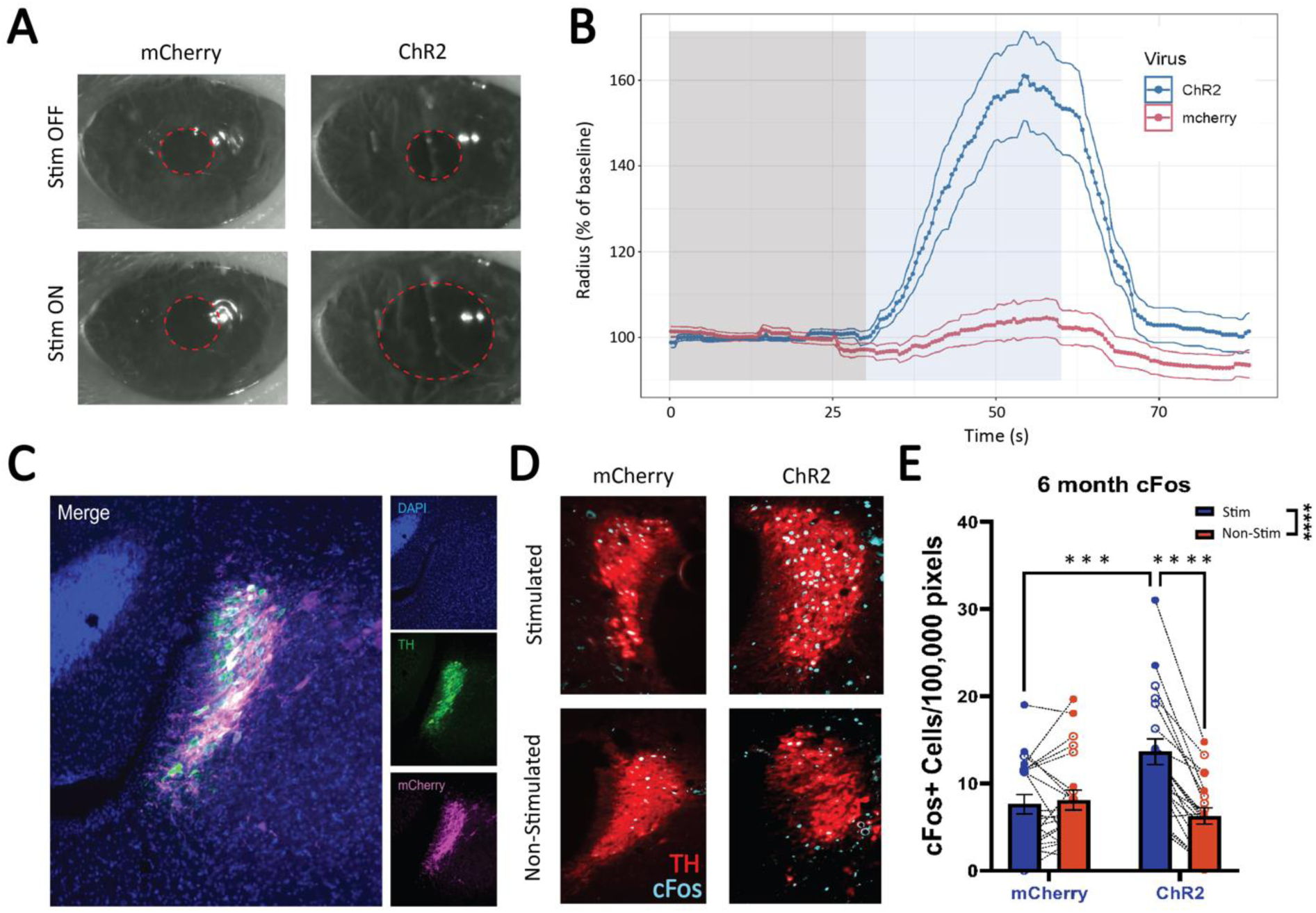
Confirmation of LC optogenetic stimulation via pupillometry and immunohistochemistry. Viral expression and response to optogenetic stimulation in the LC was verified using a combination of pupillometry and immunohistochemistry. (A) Visually notable increases in pupil diameter experienced during LC stimulation via optic fiber. (B) The change in pupil diameter was quantified and compared between mCherry and ChR2 animals using an open source pupillometry toolkit (Privitera et al., 2020). (C) Confirmation of viral expression with robust fluorescence of mCherry (pseudo-colored magenta) limited to the LC (marked by TH; green (D) c-Fos expression was used to confirm the activation of LC neurons roughly 90 min after 5 Hz constant optogenetic stimulation. Shown is a representative image of an mCherry and a ChR2 animal and the differences in c-Fos expression (teal) between the stimulated and non-stimulated hemispheres of the LC (TH; red). (E) c-Fos counts were quantified for both the mCherry and ChR2 animals at 6 months.

### 2.3 fMRI Data Acquisition

On the day of MRI scanning, rats were initially induced with 5% isoflurane for 5 minutes in combination with a 2:3 ratio of O_2_ and medical grade air. After initial induction with isoflurane, animals were maintained under 2-3% isoflurane for the remainder of handling and cradle preparations. Rats were intubated and administered the paralytic pancuronium dibromide (1.5 mg/kg/hr, s.c.; Selleck) immediately after intubation. Through the tracheal tube, breathing of the rats was maintained at 1.6 Hz using a ventilator. Animals were then placed in a custom-made MRI cradle where a custom-made transmit-receiver surface coil was placed over the brain and an optic fiber patch cord was connected to the optic ferrule. Animals were then head-fixed with ear bars and their teeth secured in a bite bar to limit head motion during scanning. To prevent the escape of light during stimulation, a black marker was used to ‘blackout’ the headcaps and tape was placed over the connection between the optic fiber and the optic ferrule implant. Eye ointment was placed over both eyes prior to scanning and isoflurane was reduced to 1.3% and maintained at this level for the rest of the scanning session. Heart rate and blood oxygen levels of the animals were monitored throughout the scanning session using a pulse oximeter that was connected to the hindpaw of the animal. Temperature of the rat during scanning was maintained at 37 ± 0.5 °C using a heated water bath system.

All fMRI scans were acquired using a 9.4T Bruker Biospec MRI scanner with a 20 cm horizontal bore. Scans were 10 minutes in length and were gradient-echo echo-planar imaging (EPI) with partial Fourier encoding with a factor of 1.4. The following parameters were used to acquire the images: isotropic voxel size = 0.5 mm^3^, matrix size = 70 x 70, slice number = 24, TE = 15 ms, TR = 1250 ms, flip angle = 90°, bandwidth = 216.45 kHz. All scans were phase-locked so that each acquisition was set to occur at a multiple of the animal’s respiratory rate. In this study, the repetition time of 1250 ms (0.8 Hz) was employed to acquire images at the frequency of every other breath of the 1.6 Hz respiration rate. This practice was adopted to ensure that each image slice was obtained at the same point in the animal’s respiratory cycle and therefore limit motion-induced artifacts as a result of chest cavity movements during respiration (Pan et al., 2020). A single volume reversed blip EPI image with the same parameters was acquired before each 10-minute scan and used for topup correction (Andersson et al., 2003; Smith et al., 2004). Saturation bands were used with all functional EPI scans to minimize signal from outside of the brain and each scan was preceded by 10 dummy scans for system calibration purposes.

Four different levels of LC stimulation were incorporated into this study: baseline (i.e., no stimulation), 2 Hz, 5 Hz, and a 15 Hz phasic stimulation (comprised of 3 pulses of light at 15 Hz every 10 seconds). These frequencies were chosen to mimic the distinct firing modes that have been observed to occur naturally in the LC: low tonic, high tonic, and phasic burst firing (Aston-Jones & Bloom, 1981; Aston-Jones & Cohen, 2005; Berridge & Waterhouse, 2003; Carter et al., 2010; McCall et al., 2015; Prokopiou et al., 2022; Valentino & Foote, 1988). Each scanning session consisted of a total of eight scans that were presented in a pseudo-randomized order: two baseline scans and two scans each at 2 Hz, 5 Hz, and 15 Hz phasic stimulation (**Figure 1**). For the scans in which 2 Hz and 5 Hz stimulation was presented, one of the two scans was acquired with constant stimulation at that frequency for the entirety of the 10 minutes, while a block design of stimulation (10 seconds of stimulation and 20 seconds of rest) was employed for the other scan. The block stimulation scans were not analyzed in the current study and will not be mentioned further. Each session started with a baseline scan, a 5 Hz constant scan was employed as the fourth scan and always followed by the second baseline scan; the placement of all other stimulation levels was randomized for each session. Placement of the 5 Hz constant stimulation scan was maintained to ensure optimal expression of c-Fos during staining approximately 90 minutes later (see section 2.5 Tissue Preparation and Immunohistochemistry) (Tillage et al., 2020, 2021). All optogenetic stimulation was done with 60mW of laser power and 10 ms-long pulses. Three minutes were allotted between scans to allow for norepinephrine levels to return to baseline post-stimulation.

### 2.4 Preprocessing

Preprocessing of fMRI data was done using the *Rodent Whole-Brain fMRI Data Processing Toolbox* (Xu, Zhang, et al., 2023). Through use of this toolbox, the following preprocessing steps were performed: slice time correction, motion correction, topup correction, nuisance regression (constant, linear, and quadratic trends as well as 6 motion regressors), normalization, bandpass filtering (0.01–0.1 Hz), spatial smoothing (FWHM = 0.5 mm), and registration to and seed extraction from the functional SIGMA-Wistar atlas (Barrière et al., 2019).

### 2.5 Tissue Preparation and Immunohistochemistry

After scanning was complete, the rats were removed from the ventilator and perfused immediately with 0.1 M kPBS followed by 4% paraformaldehyde. After brains were extracted, they were placed in 4% paraformaldehyde and stored overnight before being moved to 30% sucrose for a minimum of 3 days before slicing. Brains were sliced at 30 um at the level of the LC. Sections were either mounted on Colorfrost® Plus slides (Erie Scientific) to be counterstained with neutral red to check for optic ferrule placement (Kelberman et al., 2023) or stored in cryoprotectant until fluorescence staining.

Free floating sections were washed 3 x 5 minutes in 1x PBS and then incubated for 1 hour in blocking buffer (5% normal goat serum, 3% bovine serum albumin in 0.1% PBST). Sections were then incubated in chicken anti-tyrosine hydroxylase (TH) (1:1000; abcam ab76442) and rabbit anti-c-Fos (1:3000; abcam ab190289) for 48 hours at 4°C. Sections were then washed 3 x 5 minutes in 1x PBS and incubated for 2 hours with the corresponding secondary antibodies goat anti-rabbit 488 (1:500; Fisher Scientific A11008) and goat anti-chicken 633 (1:500; Thermo Fisher Scientific A-21103). Then, sections were washed again 3 x 5 minutes and mounted on slides, dried, and coverslipped. A separated set of LC sections were stained using the same protocols for TH and rabbit anti-DsRed (1:1000; Fisher Scientific NC9580775), along with the appropriate secondary goat anti-chicken 488 (1:500; abcam ab150169) and goat anti-rabbit 568 (1:500; Thermo Fisher Scientific A-11011) antibodies. These sections were used to visually confirm viral expression within the LC.

A Leica DM6000B epifluorescent upright microscope was used to image the sections stained for c-Fos at 10x. Using ImageJ, TH was used as a marker to manually outline the LC. A common Otsu threshold was set and despeckling was performed. Then, the number of c-Fos labelled cells was counted using standard criteria (pixel size: 100-infinity and circularity: 0.7-1.0). A two-way repeated measures ANOVA was used to assess differences in c-Fos expression across groups considering virus (ChR2-mcherry and mCherry alone) and hemisphere (stimulated and non-stimulated) as factors.

### 2.6 Data Analysis

#### 2.6.1 Spatial Distribution of the Global Signal

In order to investigate how LC stimulation affects the spatial aspect of the global signal, the regional distribution of the global signal was assessed. The global signal was calculated by extracting all timecourses within the brain region and averaging them into a single timecourse. The global signals for each scan were then z-scored by subtracting the mean and dividing by the standard deviation. The scans and global signals for each group were concatenated and the timecourse of each voxel within the brain was correlated to the global signal via a Pearson’s linear correlation. The map of correlation values of each voxel to the global signal are displayed by group. For this analysis, we examined the relative correlation with the global signal, which results in a spatial map, rather than QPP1, which results in a spatiotemporal pattern, to facilitate comparison to prior studies of the global signal. For significance testing involving the cingulate cortex, the correlation values for the three parcellations that comprised the cingulate cortex in the SIGMA-Wistar functional atlas (Cingulate Cortex 1, Cingulate Cortex 2, and Cingulate Cortex 3) were averaged to represent the activity of the cingulate cortex and a two-way ANOVA was performed considering virus (ChR2-mcherry and mCherry alone) and stimulation level (baseline, 2 Hz, 5 Hz, and 15 Hz phasic) as factors. All analysis was performed in MATLAB using custom scripts.

#### 2.6.2 QPP Analysis

QPP2 was obtained using a repeating pattern detecting algorithm that was initially reported in (Majeed et al., 2011) and later modified in (Yousefi et al., 2018). This algorithm starts by taking a template, comprised of a number of volumes within the scan that is dictated by the user-defined window length, and uses a sliding window approach to correlate this template to each timepoint of the acquired scan. A threshold is set for the correlation timecourse and at all points at which the correlation between the template and the scan is higher than the threshold, the volumes are taken and averaged to create a new template. This process is repeated until the template does not change after two iterations (cc > 0.9999). The global signal was regressed from each scan prior to applying the algorithm to ensure detection of QPP2 for comparison with prior studies (Yousefi et al., 2018). For group QPP analysis, the algorithm was run on the concatenation of all scans within said group. To assess the frequency of QPP2 occurrence for each group, a histogram of the final sliding template correlation (STC) timecourse for each group was created. STC values above the standard threshold of 0.2 were considered to be QPP events. To evaluate the pattern of the QPP2 itself, the final template provided by the algorithm was observed and compared between groups. To evaluate statistical significance, the percentage of STC values above the 0.2 threshold and a measure of cingulate cortex activity in the QPP were assessed. For the latter, the activity of the cingulate cortex (defined as the average of the three cingulate cortex parcellations defined by the SIGMA-Wistar functional atlas) during periods of detected QPP activity was averaged for each scan and the difference between the maximum and minimum levels of activity were measured. In both instances a two-way ANOVA was performed considering virus (ChR2-mcherry and mCherry alone) and stimulation level (baseline, 2 Hz, 5 Hz, and 15 Hz phasic) as factors. All QPP analyses were performed in MATLAB using custom scripts. A similar version of the scripts used in this study is available in (Xu, Yousefi, et al., 2023).

#### 2.6.3 Complex Principal Component Analysis (CPCA)

To further investigate the incidence of these spatiotemporal patterns we performed complex principal component analysis (CPCA). CPCA is able to simultaneously detect multiple spatiotemporal structures within BOLD signals, including QPP1 and QPP2 (Bolt et al., 2022). Specifically, CPCA, applied to complex Hilbert-transformed BOLD timecourses through PCA, assesses both the magnitude and phase-lag differences among brain structures. This analysis transforms BOLD signals into real (magnitude) and imaginary (phase) components, with the principal components retaining these attributes. We conducted CPCA using a Python script that was reported in (Bolt et al., 2022) and is available at https://github.com/tsb46/complex_pca. Custom MATLAB scripts were then used to evaluate the incidence of the first three principal components. Incidence of a principal component at any given timepoint was defined by which of the first three components explained the most variance in the scan at that timepoint. A two-way ANOVA (performed in MATLAB) was used to assess differences in principal component incidence across groups considering virus (ChR2-mcherry and mCherry alone) and stimulation level as factors.

## 3. RESULTS

### 3.1 Multimodal Confirmation of Optogenetic LC Stimulation

Taking advantage of the link between LC activity and pupil dilation, we used pupillometry to confirm accurate placement of the optic ferrule implant and expression of the virus (**Figure 2A**). We observed robust dilation of the ipsilateral pupil in ChR2 expressing animals during optogenetic stimulation, while stimulation-driven pupil dilation was not observed in animals expressing the mCherry control virus. Pupil dilation was quantified and compared between all mCherry and ChR2 animals (Privitera et al., 2020) (**Figure 2B**). Post-mortem staining for mCherry confirmed robust expression of the virus within the LC (**Figure 2C**). Further supporting LC activation in response to optogenetic stimulation, staining for the immediate early gene c-Fos revealed strong activation of the stimulated hemisphere only in the ChR2-expressing animals (**Figure 2D**). A two-way repeated measures ANOVA revealed no main effect of virus (F_(1,39)_ = 1.83, p = 0.18), but there was a main effect of hemisphere (F_(1, 39)_ = 29.03, p < 0.0001), subject (F_(39, 39)_ = 5.83, p < 0.0001), and a virus x hemisphere interaction (F_(1, 39)_ = 37.51, p < 0.0001). Tukey’s post-hoc tests revealed significantly elevated c-Fos in the stimulated hemisphere of ChR2 animals compared to those expressing mCherry alone (p = 0.0006) and there was a similar outcome when comparing the stimulated to non-stimulated hemisphere within ChR2 expressing animals(p < 0.0001) (**Figure 2E**).

### 3.2 Spatial Distribution of the Global Signal

To determine whether LC stimulation affects the level of contribution of different brain regions to the global signal, we assessed the spatial distribution of the global signal via a voxel-wise correlation analysis (**Figure 3**). In the mCherry control group across all levels of stimulation, higher correlation to the global signal was seen along the midline and bilaterally in posterior regions of the brain (**Figure 3A**). This finding replicates previous work that investigated the spatial distribution of the global signal in a noise-controlled cohort of rats (Anumba et al., 2023). While the same general pattern was observed in the ChR2 stimulated rats, the correlation values were widely elevated throughout the brain (**Figure 3B**). Notably, this increase was disproportionately higher in the central anterior region, which likely represents the cingulate cortex (Anumba et al., 2023), during 2 Hz and 5 Hz tonic stimulation of the LC. To further investigate this, a two-way ANOVA was performed on the correlation between the activity of the cingulate cortex and the global signal. While there were no main effects of stimulation pattern on cingulate cortex contribution to the global signal (F_(3,66)_ = 0.4204, p = 0.739), there was a main effect of virus (F_(1,66)_ = 5.836, p = 0.0185). In general, animals expressing ChR2 in the LC had stronger correlation between the activity of the cingulate cortex and the global signal. There was no additional effect based on the interaction between virus and stimulation pattern on cingulate cortex contribution to the global signal (F_(3,66)_ = 0.7455, p = 0.5288).

**Figure 3:**
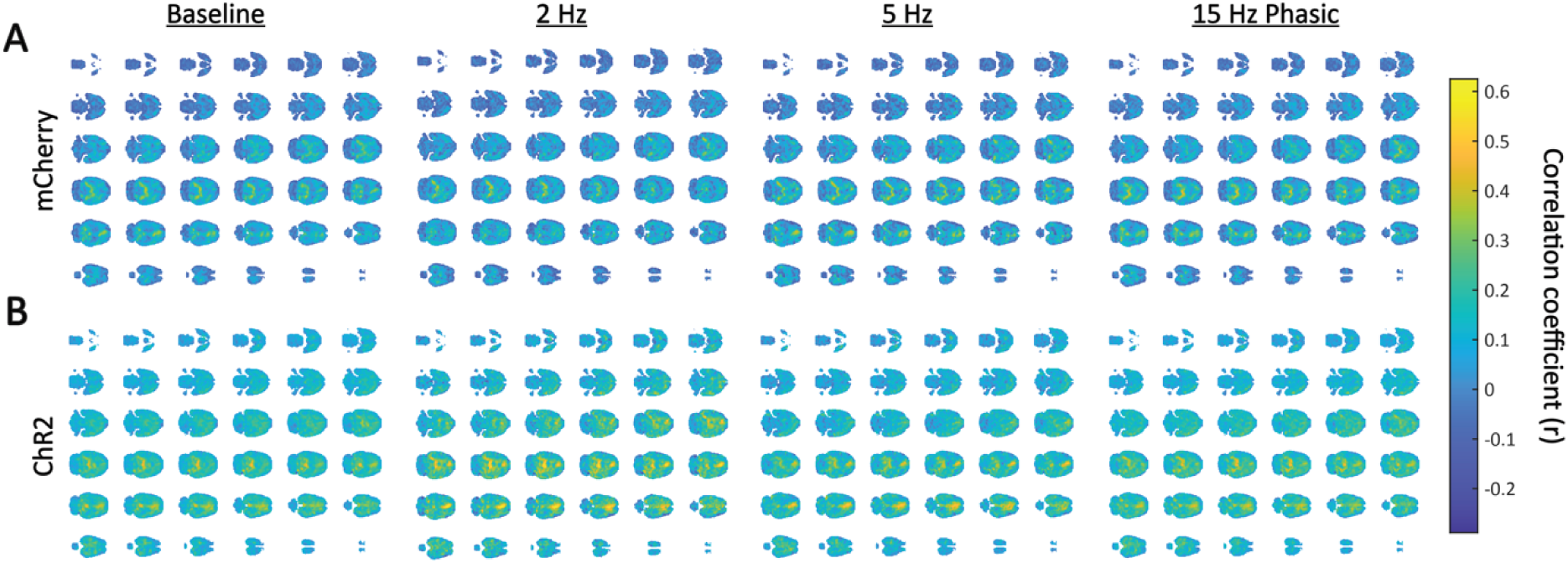
Spatial distribution of the global signal during LC stimulation. (A) The spatial distribution of the global signal in the mCherry control animals at different levels of LC stimulation. Correlation is shown in Pearson’s correlation coefficients (r). The highest correlation to the global signal was found along the midline in anterior cortical structures as well as bilaterally in posterior parts of the brain, replicating previous results (Anumba et al., 2023). (B) The spatial distribution of the global signal in ChR2 stimulated animals at different levels of LC stimulation. The same pattern of highly correlated structures as seen in the control animals was observed here although with higher correlation seen throughout the whole brain. When compared to baseline, correlation in the bilateral posterior region was stronger during 2 Hz LC stimulation and correlation in the anterior medial structure was stronger during 2 Hz and 5 Hz stimulation of the LC.

### 3.3 QPP2 Analysis

A pattern finding algorithm was used to detect the occurrence of QPP2 in this dataset (see Methods for details). The STC values of the final QPP template with the scans of each group are shown plotted as a histogram (**Figure 4A**). The threshold for a QPP event was set to 0.2 and any timepoints at which correlation was above this threshold were considered QPP events. The distributions of correlation values for all scan groups showed a near complete overlap between the mCherry and ChR2 animals at baseline. However, for the distributions during the 2 Hz and 5 Hz stimulation levels, a higher percentage of correlation values above the absolute value of the 0.2 threshold were observed in the ChR2 stimulated animals than in the mCherry controls. This indicates that more QPP events were detected in the ChR2 animals during 2 Hz and 5 Hz stimulation when compared to the mCherry controls. During 15 Hz phasic stimulation, there was more overlap between mCherry and ChR2 animals above the threshold, although this overlap was not as complete as it is for the baseline scans. To evaluate these trends, a two-way ANOVA was performed on the percentage of STC values above the 0.2 threshold. However, we found no effect of stimulation pattern (F_(3,66)_ = 0.7457, p = 0.5287), virus (F_(1,66)_ = 1.933, p = 0.1691), or a stimulation pattern x virus interaction (F_(3,66)_ = 0.962, p = 0.416) on the percentage of STC values above the 0.2 threshold.

**Figure 4:**
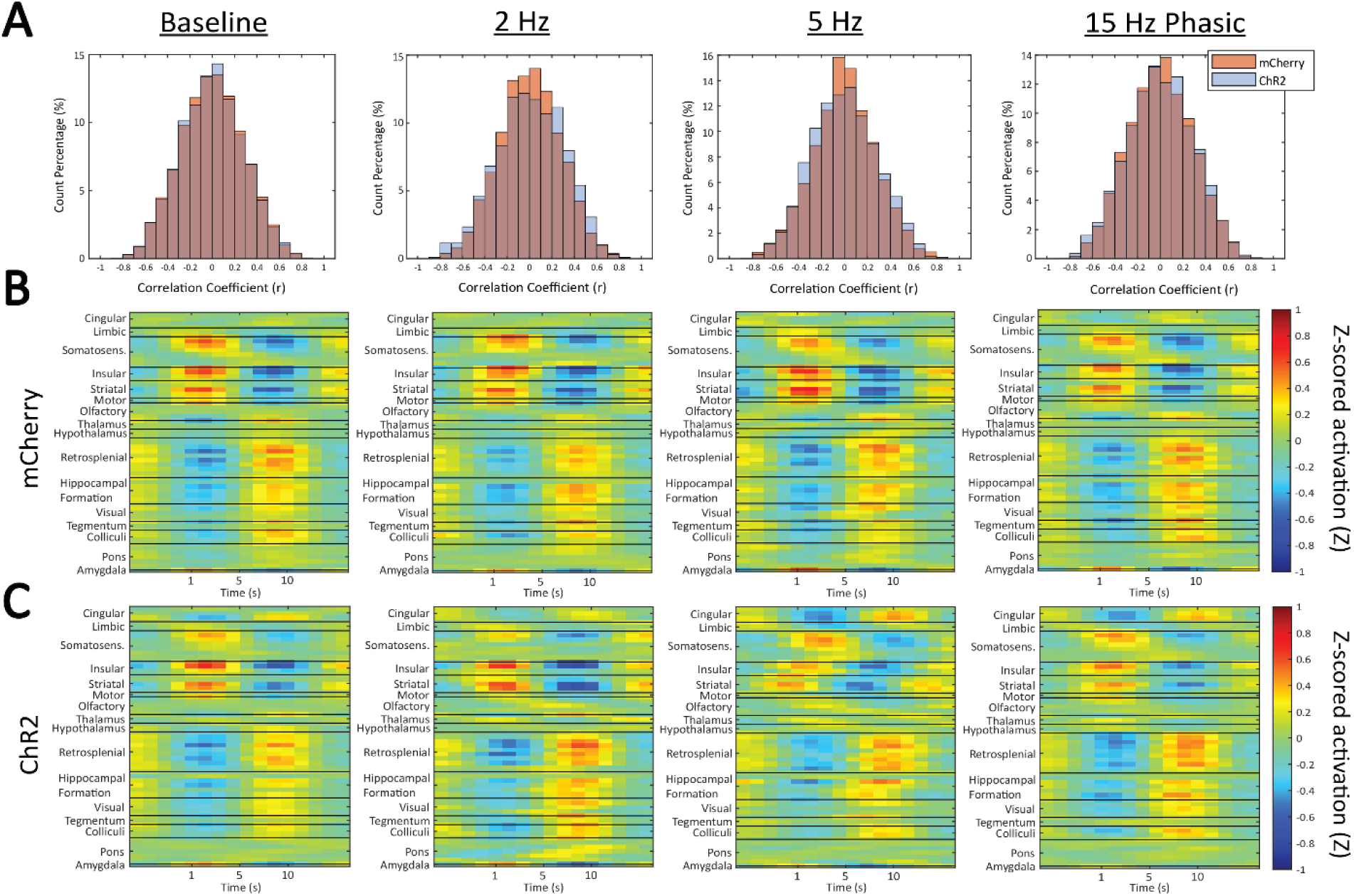
QPP detection and templates. (A) STC histograms for mCherry (orange) and ChR2 (blue) animals where values over 0.2 were considered QPP events. Very little difference was shown between the two animal groups; however, a slightly higher percentage of counts were above the 0.2 threshold for the ChR2 animals during 2 Hz and 5 Hz stimulation. (B) QPP templates, or regional activity within the QPP, for mCherry animals at each stimulation level. (C) QPP templates, or regional activity within the QPP, for ChR2 animals at each stimulation level. Some minor differences in regional activity were seen during 5 Hz tonic and 15 Hz phasic LC stimulation.

The QPP template generated for each scan group was also compared across groups to assess the involved activity of each brain region within the QPP (**Figure 4B-C**). The templates show the waves of activation and deactivation that were exhibited by distinct regions and that characterize the QPP. Notably, certain regions exhibited activity that was in phase with each other and distinctly out of phase with other regions, whereas other regions showed very little oscillatory activity in the pattern at all. The somatosensory, insular, striatal, motor, and amygdala regions all exhibited similar activity that was almost exactly anticorrelated to the activity of the retrosplenial cortex, hippocampal formation, and certain deep brain structures. Other cortical regions, such as the olfactory network and cingulate cortex, and sub-cortical regions such as the thalamus, hypothalamus, and the pons, contributed very little to the dynamic pattern. Importantly, the cingulate cortex showed a much stronger involvement in the QPP of ChR2 animals under 5 Hz tonic and 15 Hz phasic LC stimulation compared to any of the control animals or at other stimulation levels in the same group. This cingulate activity, when present, moved in phase with the retrosplenial and hippocampal regions. Once again, a two-way ANOVA revealed no effect of stimulation pattern (F_(3,66)_ = 0.8452, p = 0.4741), virus (F_(1,66)_ = 0.1287, p = 0.721), or a stimulation pattern x virus interaction (F_(3,66)_ = 0.3243, p = 0.8087) on the degree to which the cingulate was involved in the QPP. However, we did observe an apparent phase shift in the activity of the insular and striatal regions when compared to that of the somatosensory network during 5 Hz stimulation in the ChR2 animals.

### 3.4 Complex Principal Component Analysis (CPCA)

Using the unique feature of CPCA to evaluate regional differences in time-lag activity, we performed this analysis as a complementary way to observe the spatiotemporal dynamics in the dataset. Previous work in humans revealed the first three principal components of CPCA as common spatiotemporal patterns of activity often observed in the brain (Bolt et al., 2022). Whether detected with the QPP algorithm or CPCA, the first component represents the BOLD global signal, and the second component captures the observed anti-correlation between the DMN and TPN. For this reason, CPCA results in this study were confined to the first three principal components (Meyer-Baese et al., 2024). In the interest of continuity, in this section we will refer to the first three principal components as QPP1, QPP2, and QPP3, to remain consistent with the nomenclature used for the QPP algorithm.

The effects of LC stimulation on the incidence of each QPP were analyzed by comparing the percentage of scan time that was dominated by each component for the mCherry control group and the ChR2 stimulation group (**Figure 5**). As expected, across groups and stimulation levels, QPP1 was responsible for a majority of the variance explained (∼70%). Incidence of QPP1 was slightly higher for all ChR2 animals as indicated by a two-way ANOVA that reported no effect of stimulation level (F_(3, 66)_ = 0.68, p = 0.5691), but a significant effect of the virus (F_(1, 66)_ = 6.19, p = 0.0154). There was no additional effect on QPP1 incidence based on the interaction between virus and stimulation pattern (F_(3,66)_ = 0.29, p = 0.8329). Generally, animals expressing ChR2, regardless of stimulation pattern, spent a higher proportion of time in QPP1 compared to mCherry-expressing animals. The overall incidence of QPP2 was much lower than was observed for QPP1, however incidence of QPP2 was consistently lower in ChR2 animals when compared to mCherry control animals. A two-way ANOVA revealed no effect of stimulation level (F_(3, 66)_ = 0.39, p = 0.7578), but there was a significant effect of the virus (F_(1, 66)_ = 6.42, p = 0.0137). There was no additional effect on QPP2 incidence based on the interaction between virus and stimulation pattern (F_(3,66)_ = 0.8, p = 0.501). Regardless of stimulation level, ChR2-expressing animals generally spent a lower proportion of time in QPP2 compared to mCherry animals. The overall incidence of QPP3 was the lowest of the three components with no consistent trend observed across stimulation level or virus. A two-way ANOVA revealed no significant effect of stimulation level (F_(3, 66)_ = 0.82, p = 0.4892) or virus (F_(1, 66)_ = 0.89, p = 0.3486). The timing and incidence of each component during the scan was also assessed at the individual level, however no trend was observed as a result of LC stimulation (**Figure S2**).

**Figure 5:**
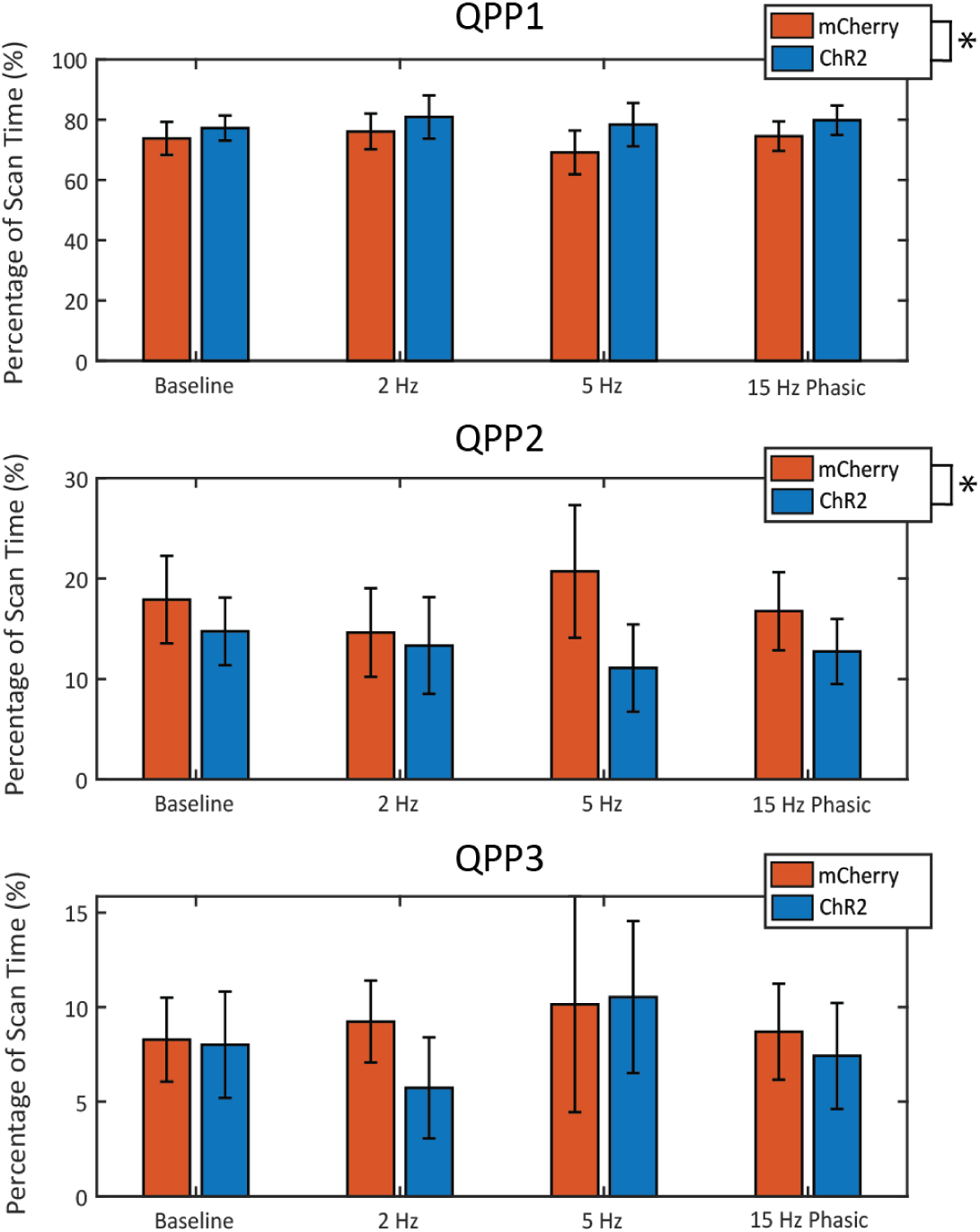
Incidence percentage of the first three QPPs during LC stimulation. The percentage of scan time for which each of the first three QPPs was dominant is displayed in this figure. The differences between the control animals (orange) and the stimulated animals (blue) were compared for each level of LC stimulation. Incidence for QPP1 across all groups was much higher than it was for QPP2 and QPP3. A two-way ANOVA revealed significance on account of virus for the first two components, however post-hoc analysis showed no further significance at the level of stimulation. Error bars represent the mean +/− SEM.

## 4. DISCUSSION

Both the BOLD global signal and the anticorrelation between DMN and TPN are associated with arousal and measures of vigilance (Pisauro et al., 2016; Wong et al., 2012, 2013, 2016). The LC plays critical role in arousal, exerting its neuromodulatory influence throughout the brain (Aston-Jones & Cohen, 2005; Berridge & Waterhouse, 2003; Poe et al., 2020). Through the use of optogenetic-fMRI, we have shown that stimulation of the LC at different frequencies results in relatively small changes in correlation to global signal that can be significant in some regions of the brain, and that ChR2-expressing rats exhibit higher ratios of QPP1 to QPP2 than mCherry control rats.

### 4.1 LC Stimulation and the Global Signal

The global signal exhibits an anticorrelated relationship with arousal and vigilance measures across species (Chang et al., 2016; Pisauro et al., 2016; Wong et al., 2013, 2016). Given this relationship with arousal, we were curious about how the global signal would be affected by stimulation of the LC. With respect to the spatial distribution of the global signal, LC stimulation did indeed appear to result regionally specific changes. First, the primary layout of correlation to the global signal that was observed in both the mCherry and ChR2 animals reflected that reported in (Anumba et al., 2023), where the midline structure, which likely includes the cingulate cortex, showed the highest correlation to the global signal. Interestingly, stimulation of the LC in the ChR2 animals did not equally increase correlation to the global signal throughout the brain, but instead resulted in higher correlation from regions that previously showed elevated global signal correlation (**Figure 3**). In (Anumba et al., 2023), the DMN was shown to contribute significantly to the global signal in rats, and when matching regions of high global signal correlation to the rat DMN as reported in (Lu et al., 2012), the strongly correlated midline structures were comprised of the cingulate cortex and parts of the retrosplenial cortex. Studies in humans have also shown that certain DMN nodes, including the posterior cingulate cortex for which the retrosplenial cortex is an ortholog, have shown both higher activity than the global signal (Raichle et al., 2001) and a higher amplitude of ultra-low frequency fluctuations than the global signal (Yu-Feng et al., 2007). The anterior cingulate cortex (ACC), which corresponds to the cingulate cortex in rodents, is involved in a wide array of high-order cognitive functions such as the regulation of emotions, decision making, and learning (Apps et al., 2016; Stevens et al., 2011). A human brain mapping study that considered brainstem structures showed that the ACC had the strongest summed functional connectivity to all brainstem nuclei, making it a primary hub for cortical-brainstem functional connectivity (Hansen et al., 2023). Stimulation of alpha-2a adrenergic receptors in non-human primates was reported to increase the encoding of negative prediction errors in the ACC, linking the noradrenergic system to the ACC’s involvement in learning (Hassani & Womelsdorf, 2023). Our lab has also shown the importance of reciprocal connections between the ACC and LC in affective responses to contextual novelty (Lustberg, Iannitelli, et al., 2020; Lustberg, Tillage, et al., 2020). Therefore, we asked whether LC stimulation resulted in increased correlation of the cingulate cortex to the global signal. The largest effects were observed during tonic 2 Hz and 5 Hz stimulation of the LC, which are associated with generalized awake states and stress, respectively (Aston-Jones & Bloom, 1981; Aston-Jones & Cohen, 2005; Berridge & Waterhouse, 2003; Carter et al., 2010; McCall et al., 2015; Morris et al., 2020; Prokopiou et al., 2022; Valentino & Foote, 1988). Importantly, the increase in correlation of the cingulate cortex to the global signal was apparent within ChR2 animals compared to mCherry animals as a whole, rather than based on specific stimulation patterns. We hypothesize that this may be due to the cumulative and lasting effects of LC stimulation over the entire imaging period, which may have influenced global signal even during baseline scans. The lack of significant effects on the basis of stimulation pattern indicates that observed trends, such as the increase in cingulate cortex contribution to the global signal, may be small in nature. It is also important to note that the group sizes for some stimulation levels, especially for the ChR2 animals, were relatively small, making it possible that more power afforded by bigger group sizes could uncover a significant effect.

### 4.2 LC Stimulation and QPP2

QPP2 was detected using the robust pattern-detection algorithm reported in (Yousefi et al., 2018), although there was very little change in the detection rate as a result of LC stimulation (**Figure 4A**). However, the histograms of STC values during different levels of stimulation revealed a slightly higher, but non-significant, percentage of correlation values above the 0.2 threshold in the ChR2 animals during 2 Hz and 5 Hz LC stimulation. It is interesting to compare these findings with a preliminary study in which either elevating extracellular NE levels with the NE transporter inhibitor atomoxetine or depleting NE using the LC-specific neurotoxin DSP4 disrupted the generation of QPPs (Abbas et al., 2018). By contrast, in the current study we found that tonic stimulation of the LC *increases* the detection of QPPs. An important distinction between these two studies is that atomoxetine and DSP4 induce dramatic changes in LC-NE signaling, whereas optogenetic stimulation triggers more subtle effects that mimic physiological function. One interpretation consistent with the “adaptive gain theory” and inverted U dose-response nature of LC function (Aston-Jones & Cohen, 2005) is that “optimal” LC-NE transmission promotes QPP2, while pathological changes to LC circuits associated with neuropsychiatric and neurodegenerative diseases intrude on these brain states (Iannitelli & Weinshenker, 2023; Kelberman et al., 2020; Poe et al., 2020; Weinshenker, 2018; Weinshenker & Holmes, 2016).

The relative regions involved in the detected QPP as well as their activity during the pattern were also observed. The foundational structure of the QPP was very similar across groups and stimulation levels, with certain regions exhibiting stronger activity within the pattern and those regions belonging to one of two groups of structures whose activity is anticorrelated (**Figure 4B-C**). In particular, the somatosensory, insular, and striatal networks move in phase with each other whereas the retrosplenial cortex, hippocampus, visual network, and a few midbrain structures were in phase with each other. These groupings of regions within the QPP largely reflect what has previously been reported for QPPs in rats (Majeed et al., 2011; van den Berg et al., 2022). Interestingly, both of these studies show strong involvement of the cingulate cortex in the QPP at rest, whereas in our study strong activity from the cingulate cortex is only observed at the group level during stimulation of the LC at 5 Hz tonic and 15 Hz phasic levels. Under these conditions, the activity of the cingulate cortex is in phase with the retrosplenial cortex and hippocampus both of which, in addition to the cingulate cortex, have been defined as part of the rat DMN (Lu et al., 2012). LC firing at 5 Hz tonic and 15 Hz phasic levels is associated with anxiety and focused attention, respectively (Berridge & Waterhouse, 2003; Carter et al., 2010; McCall et al., 2015). These results indicate that LC activity during states of high arousal, regardless of valence, could lead to an increase in activity of the cingulate cortex in rats, especially in the context of slow spatiotemporal patterns like QPPs. Importantly, the effect of stimulation on increases in cingulate activity in the QPP were not statistically significant, indicating that this effect also may be small in nature. That being said, it known that the pattern-detecting algorithm used to detect the QPPs is more robust at the group level (i.e., with more data) and that results on the individual level are not necessarily indicative of the group level results (H. Watters et al., 2024). It should also be noted that during 5 Hz LC stimulation there was a slight phase shift in the activity of the insular and striatal networks, meaning that their peaks in activity occurred slightly before those of the other structures. Though this effect has not previously been reported, it is a point of interest that could be further investigated.

### 4.3 CPCA Findings

We used CPCA to assess whether LC stimulation had any effect on the presentation and occurrence of the first three principal components (QPP1, QPP2, QPP3) in these rats. A recent study showed that the majority of large-scale spatiotemporal signals that have been detected with fMRI can be explained by the first three principal components, or QPPs, of activity as calculated through CPCA (Bolt et al., 2022). Specifically, they showed that QPP1 was reflective of the global signal and that QPP2 was indicative of what is defined as the anticorrelated QPP. Moreover, recent work in rats showed that different anesthetic conditions alter the ratio of QPPs, and in humans, QPP1 increases at the expense of QPP2 over the course of a scan (Meyer-Baese et al., 2024).

LC stimulation appeared to have very subtle effects on the incidence of the principal components despite consistent trends being observed based on which virus animals expressed (**Figure 5**). ChR2 expression should not cause significant changes alone, and we believe this effect could be due to lasting effects of LC stimulation across scans in ChR2-expressing animals or as a result of other variables, such as anesthesia, which can produce large changes in the distribution of these QPPs (Meyer-Baese et al., 2024). Regardless, the comparison that produced the largest effect size was the incidence of QPP2 during 5 Hz stimulation (approximately 10% lower occurrence in ChR2 animals). This is worth noting because our other findings using the QPP pattern-detecting algorithm reveal that stimulation at 5 Hz induced changes in the presentation of the QPP (**Figure 4**). Given this and the fact that QPP2 is reflective of the anticorrelation between DMN and TPN (Bolt et al., 2022), it is possible that these changes during 5 Hz stimulation could be responsible either for the signal that was detected as QPP2 when using CPCA or for the reduced frequency of QPP2 occurrence that we see in these scans.

### 4.4 Potential Subtle Effects

Despite the LC having global neuromodulatory influence, the majority of the results presented here display effects that are quite subtle, highlighted by the lack of statistical significance. Moreover, observed effects were restricted to specific brain regions or structures as opposed to the whole brain. It should be noted that the effects of stimulating small nuclei using optogenetic-fMRI are usually quite small. Studies using optogenetics to stimulate the ventral posteromedial nucleus (VPM) of the thalamus and LC have reported significant yet small changes in % BOLD signal as a result of optogenetic stimulation (Chuang et al., 2023; Grimm et al., 2022). In contrast, our study did not show significant effects as a result of LC stimulation, and the effects seen were not observed on a global scale as has been reported by others (Oyarzabal et al., 2021; Zerbi et al., 2019). However, the latter may be attributed to the differences in method of LC stimulation. Correspondingly, the sensitivity of optogenetic-fMRI is thought to potentially underreport whole-brain responses when used to stimulate small areas (Lee et al., 2022).

As mentioned above, the small effects that were observed could also be explained by the Yerkes-Dodson model in which the relationship between arousal and performance is explained by an inverted-U-shaped curve, with a small range of arousal leading to optimal performance (P. A. Watters et al., 1997). This inverted-U curve has also been shown to explain the relationship between LC-NE modulation and network function and behavior (Aston-Jones & Cohen, 2005; Poe et al., 2020). When compared to extreme forms of LC-NE manipulation (Abbas et al., 2018), optogenetic modulation of the LC allows a more fine-tuned approach over a narrow range of activity that may result in more subtle effects. Additionally, although our study was designed to account for little interference of between-scan effects, a study showed that high tonic firing of the LC can reduce the excitability and NE release of subsequent phasic firing modes (Li et al., 2023). Therefore, it is possible that the length of our scans (10 min) with constant stimulation may have resulted in overall NE *depletion*, resulting in a lack of effects observed over the whole length of the scan. Reanalysis using a shorter stimulation period would be a necessary direction for future study. Given the nature of the BOLD signal and its dependence on signal from the brain vasculature, it is important to recognize the role of NE as a vasoconstrictor and how this could have resulted in competing effects to the signal captured during LC stimulation (Bekar et al., 2012). This also speaks to the complexity of the neural and hemodynamic systems involved when measuring these spatiotemporal signals with fMRI. Lastly, despite the valuable advancements that work in rodents allows (Pagani et al., 2023), it is important to note that the animals presented were scanned under anesthesia, a state which is known to affect measures of brain activity (X. Liu et al., 2013; Masamoto et al., 2009; Masamoto & Kanno, 2012), and may compete with arousal-associated manipulations such as certain levels of LC stimulation.

### 4.5 Limitations and Future Directions

One limitation of this study is the unilateral nature of our LC optogenetic stimulation. Plenty of studies have induced behavioral phenotypes utilizing unilateral optogenetic LC stimulation (Marzo et al., 2014; McCall et al., 2015; Tillage et al., 2020, 2021). However, a majority of LC fibers project ipsilaterally and some have produced unilateral effects as a result of LC stimulation (Grimm et al., 2022; Marzo et al., 2014; Proudfit & Clark, 1991; Waterhouse et al., 1983, 2022), something that this study was unable to assess due to the lack of bilateral symmetry in the functional atlas used during analysis. Therefore, more robust effects may be observed with bilateral stimulation, as has been observed previously (Zerbi et al., 2019). Furthermore, while we utilized optogenetics for the added benefits of controlling precise timing and patterns of LC stimulation, this approach is limited by light penetration through the tissue, variable viral expression, and placement of optic ferrules. All of these could impact the total number of LC neurons and location within the LC core that were responsive to optogenetic stimulation. Given the emerging appreciation of the heterogeneity of the LC (Ma et al., 2023; Poe et al., 2020; Schwarz & Luo, 2015; Uematsu et al., 2015), our manipulations could have been targeting specific downstream brain regions rather than influence brain wide networks. Follow up studies could isolate local effects of LC activity on specific brain regions by stimulating LC axon terminals. Lastly, the reported activity of certain brainstem structures should be taken cautiously due to the use of a surface coil which was unable to obtain optimal signal from structures at this depth.

## 5. CONCLUSION

In this study we combined optogenetic stimulation of the LC with whole-brain fMRI recordings to assess the effects of LC activity on large-scale spatiotemporal dynamics. Specifically, we assessed the spatial components of the BOLD global signal, the presentation of QPPs, and the principal components extracted from CPCA as a result of LC stimulation. Our findings provide evidence that LC stimulation at specific levels results in small regionally specific effects with regards to the global signal and QPPs. LC stimulation showed little change in the frequency of occurrence of CPCA principal components. These results suggest that the effects of LC stimulation are not simultaneously widespread and large in scale, but instead result in more regional effects that are small in nature. This study contributes an additional insight to our wholistic understanding of how resting-state global dynamics are affected by ongoing neuromodulation and differing brain states such as arousal.

## 6. DATA AND CODE AVAILABILITY

A MATLAB script with the code necessary to perform the reported analyses will be available via GitHub once the final version of the paper is accepted. rs-fMRI data will be available upon request.

## 7. AUTHOR CONTRIBUTIONS

NA and MK worked equally in terms of project conceptualization, data collection, data analysis, and manuscript writing. WP assisted in optogenetic-fMRI optimization, data collection, and experimentation. AM assisted with processing and analyzing pupillometry videos. XZ assisted with manuscript revisions. NX assisted with data preprocessing troubleshooting, as well as manuscript revisions and editing. DW and SK were responsible for supervision of the project, review, and editing.

## 9. ACKNOWLEDGEMENTS

The authors would like to thank the Center for Systems Imaging Core and the Cloning and Viral Vector Cores at Emory University for enabling use of the 9.4 T MRI scanner and for producing the viruses used in the optogenetic experiments. We would also like to thank C. Smith, G. P. Clavijo, and L. Daley for their assistance in data collection and E. Kim for her help with pupillometry analysis.

## 10. SUPPLEMENTARY MATERIAL

**Supplementary Table 1:**
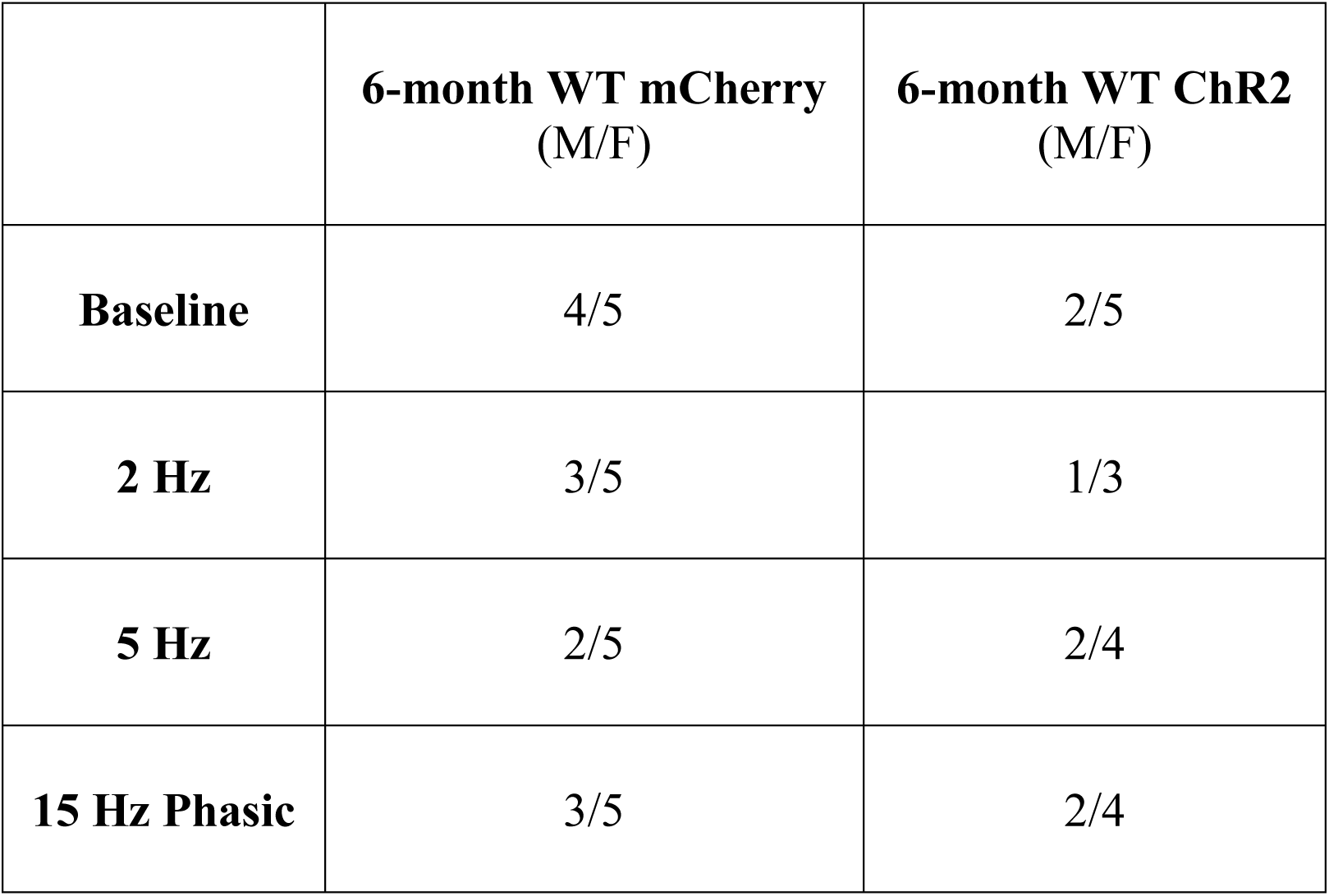
Animal Group Sizes by Sex.

### 10.1 Power Spectral Density Analysis of the Global Signal

**Figure S1:**
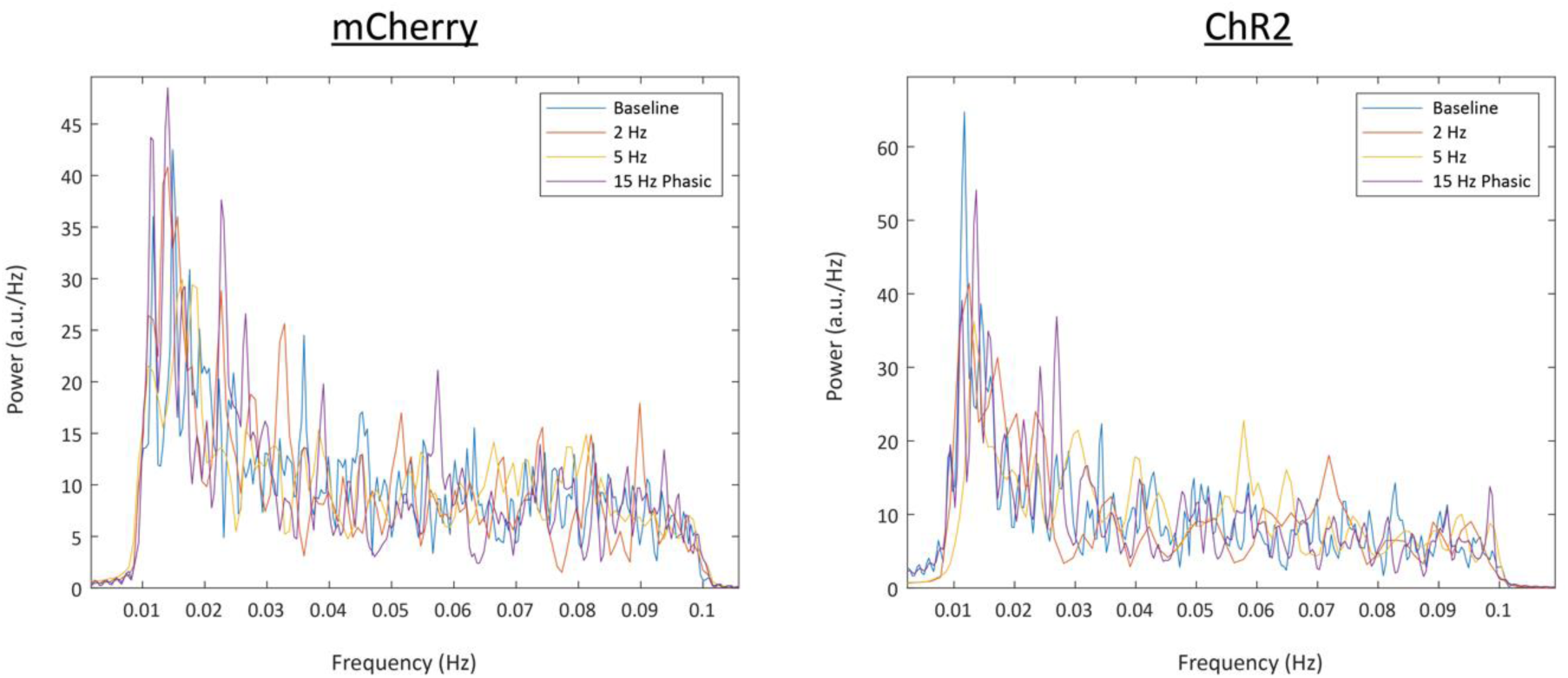
Power spectral density (PSD) estimates of the global signal. PSD estimates for the mCherry control animals (left) and the ChR2 stimulated animals (right) are displayed in this figure. All stimulation groups showed a high low-frequency peak in power with a gradual decrease in power as the frequency increased. In the ChR2 animals, the power of this low-frequency peak was higher in the baseline and 15 Hz phasic scans when compared to the control animals. However, no other strong differences were observed between the two groups of animals.

The frequency of the global signal was observed by way of a power spectral density (PSD) analysis. PSD estimates were obtained in MATLAB using Welch’s method with a Hamming window and 50% overlap. The global signals for each group were z-scored and concatenated, after which a PSD was calculated for the concatenated signals producing the group results. A PSD estimate was created for the scans at each stimulation level for both the mCherry control and ChR2 stimulated rats (**Figure S1**). Typical for resting-state fMRI scans (Pan et al., 2013), at all stimulation levels a low frequency (< 0.02 Hz) peak in activity was observed, after which the magnitude of power proceeded to steadily taper off at higher frequencies. The magnitude of this low frequency peak was maintained around the same level in the ChR2 stimulated animals when compared to the mCherry controls, with the exception of the baseline and 15 Hz phasic scans for which the low frequency peak was slightly larger in the ChR2 animals. However, no strong notable differences were observed in the power spectra when comparing the corresponding traces for the ChR2 animals to the mCherry controls. In fact, the largest noticeable difference was observed in the baseline scans. It is important to note that this data was bandpass filtered between 0.01 – 1.0 Hz to highlight ultralow fluctuations associated with resting-state activity (Pan et al., 2013). Therefore, it is possible that higher frequencies contained in the global signal may be significantly affected by LC stimulation.

### 10.2 Individual CPCA Incidence

**Figure S2:**
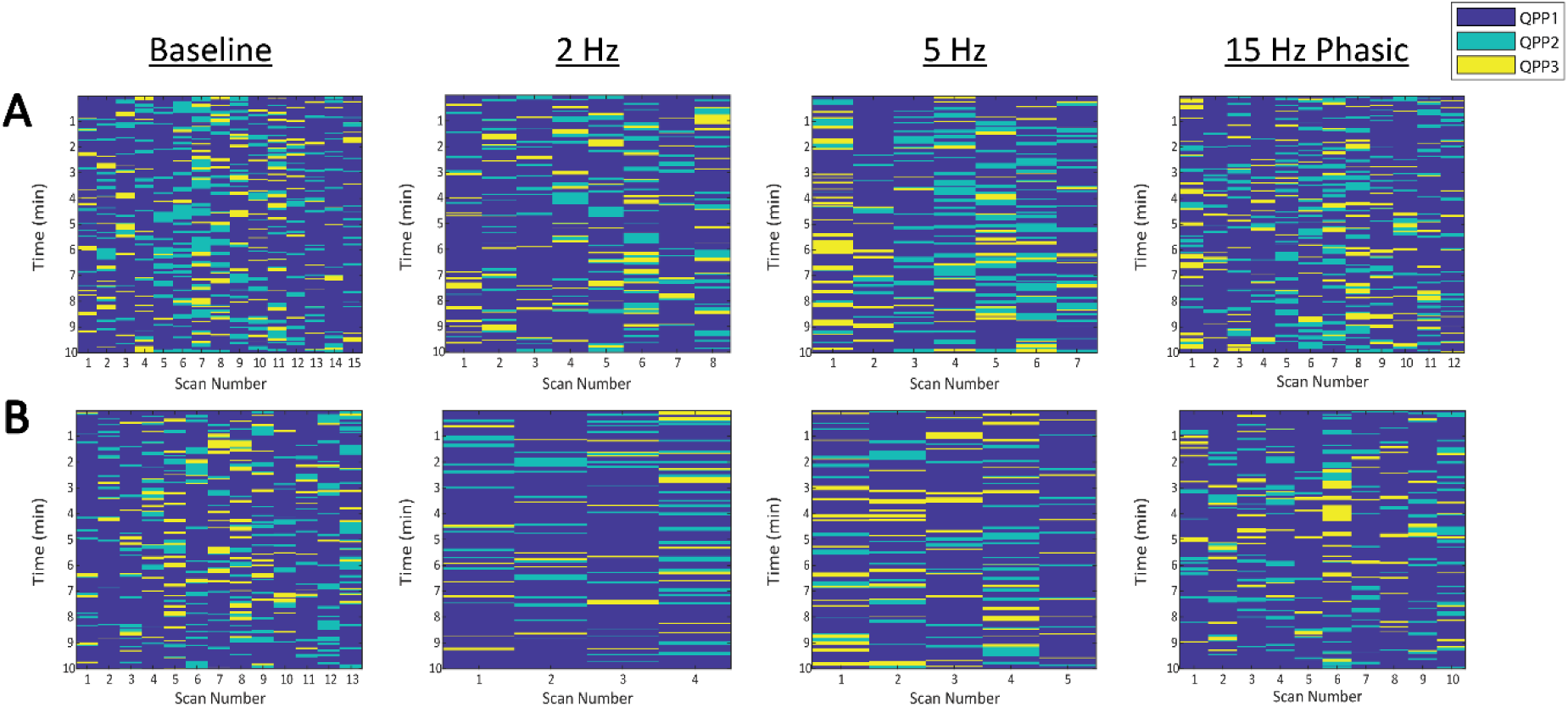
Incidence of the first three QPPs throughout each scan during LC stimulation. Each graph shows the number of scans in each group and the principal component that was dominant at each timepoint throughout the scan. The results for the mCherry control animals (A) and ChR2 stimulated animals (B) are shown. These graphs show strong individual variability in terms of timing and the temporal distribution of components across scans and across groups.

The timing of when a component was dominant throughout the scan proved to be highly varied across scans (**Figure S2**). A majority of scan time was dominated by QPP1 with short periods of QPP2 and QPP3 interspersed throughout the scan. Although overall the least amount of time was spent in QPP3, certain scans were seen to spend more time in QPP3 than others (such as scan 1 in the mCherry 5 Hz group and scan 6 in the ChR2 15 Hz phasic group). While longer periods of time (sometimes on the order of minutes) were observed to remain in QPP1, the positioning of these time periods with respect to the beginning and the end of the scan varied quite a bit across rats. No observable pattern as to the timing and sequence of the varying component states was seen based on virus or stimulation level.

## REFERNECES

1. Abbas, A., Bassil, Y., & Keilholz, S. (2019). Quasi-periodic patterns of brain activity in individuals with attention-deficit/hyperactivity disorder. NeuroImage: Clinical, 21, 101653. 10.1016/j.nicl.2019.101653

2. Abbas, A., Belloy, M., Kashyap, A., Billings, J., Nezafati, M., Schumacher, E. H., & Keilholz, S. (2019). Quasi-periodic patterns contribute to functional connectivity in the brain. NeuroImage, 191, 193–204. 10.1016/j.neuroimage.2019.01.076

3. Abbas, A., Majeed, W., Thompson, G., & Keilholz, S. (2016). Phase of quasi-periodic patterns in the brain predicts performance on psychomotor vigilance task in humans. International Society for Magnetic Resonance Iin Medicine. https://archive.ismrm.org/2016/1192.html

4. Abbas, A., Nezafati, M., Thomas, I., & Keilholz, S. (2018). Quasiperiodic Patterns in BOLD fMRI Reflect Neuromodulatory Input. International Society for Magnetic Resonance in Medicine. https://index.mirasmart.com/ISMRM2018/PDFfiles/1118.html

5. Abbott, S. B. G., Stornetta, R. L., Fortuna, M. G., Depuy, S. D., West, G. H., Harris, T. E., & Guyenet, P. G. (2009). Photostimulation of retrotrapezoid nucleus phox2b-expressing neurons in vivo produces long-lasting activation of breathing in rats. The Journal of Neuroscience: The Official Journal of the Society for Neuroscience, 29(18), 5806–5819. 10.1523/JNEUROSCI.1106-09.2009

6. Andersson, J. L. R., Skare, S., & Ashburner, J. (2003). How to correct susceptibility distortions in spin-echo echo-planar images: Application to diffusion tensor imaging. NeuroImage, 20(2), 870–888. 10.1016/S1053-8119(03)00336-7

7. Anumba, N., Maltbie, E., Pan, W.-J., LaGrow, T. J., Xu, N., & Keilholz, S. (2023). Spatial and Spectral Components of the BOLD Global Signal in Rat Resting-State Functional MRI. Magnetic Resonance in Medicine, 90(6), 2486–2499. 10.1002/mrm.29824

8. Apps, M. A. J., Rushworth, M. F. S., & Chang, S. W. C. (2016). The Anterior Cingulate Gyrus and Social Cognition: Tracking the Motivation of Others. Neuron, 90(4), 692–707. 10.1016/j.neuron.2016.04.018

9. Aston-Jones, G., & Bloom, F. E. (1981). Activity of norepinephrine-containing locus coeruleus neurons in behaving rats anticipates fluctuations in the sleep-waking cycle. The Journal of Neuroscience: The Official Journal of the Society for Neuroscience, 1(8), 876–886. 10.1523/JNEUROSCI.01-08-00876.1981

10. Aston-Jones, G., & Cohen, J. D. (2005). AN INTEGRATIVE THEORY OF LOCUS COERULEUS-NOREPINEPHRINE FUNCTION: Adaptive Gain and Optimal Performance. Annual Review of Neuroscience, 28(1), 403–450. 10.1146/annurev.neuro.28.061604.135709

11. Barrière, D. A., Magalhães, R., Novais, A., Marques, P., Selingue, E., Geffroy, F., Marques, F., Cerqueira, J., Sousa, J. C., Boumezbeur, F., Bottlaender, M., Jay, T. M., Cachia, A., Sousa, N., & Mériaux, S. (2019). The SIGMA rat brain templates and atlases for multimodal MRI data analysis and visualization. Nature Communications, 10(1), Article 1. 10.1038/s41467-019-13575-7

12. Bekar, L. K., Wei, H. S., & Nedergaard, M. (2012). The Locus Coeruleus-Norepinephrine Network Optimizes Coupling of Cerebral Blood Volume with Oxygen Demand. Journal of Cerebral Blood Flow & Metabolism, 32(12), 2135–2145. 10.1038/jcbfm.2012.115

13. Belloy, M. E., Naeyaert, M., Abbas, A., Shah, D., Vanreusel, V., van Audekerke, J., Keilholz, S. D., Keliris, G. A., Van der Linden, A., & Verhoye, M. (2018). Dynamic resting state fMRI analysis in mice reveals a set of Quasi-Periodic Patterns and illustrates their relationship with the global signal. NeuroImage, 180, 463–484. 10.1016/j.neuroimage.2018.01.075

14. Benarroch, E. E. (2018). Locus coeruleus. Cell and Tissue Research, 373(1), 221–232. 10.1007/s00441-017-2649-1

15. Berridge, C. W., & Waterhouse, B. D. (2003). The locus coeruleus–noradrenergic system: Modulation of behavioral state and state-dependent cognitive processes. Brain Research Reviews, 42(1), 33–84. 10.1016/S0165-0173(03)00143-7

16. Birn, R. M., Diamond, J. B., Smith, M. A., & Bandettini, P. A. (2006). Separating respiratory-variation-related fluctuations from neuronal-activity-related fluctuations in fMRI. NeuroImage, 31(4), 1536–1548. 10.1016/j.neuroimage.2006.02.048

17. Bolt, T., Nomi, J. S., Bzdok, D., Salas, J. A., Chang, C., Thomas Yeo, B. T., Uddin, L. Q., & Keilholz, S. D. (2022). A parsimonious description of global functional brain organization in three spatiotemporal patterns. Nature Neuroscience, 25(8), Article 8. 10.1038/s41593-022-01118-1

18. Carter, M. E., Yizhar, O., Chikahisa, S., Nguyen, H., Adamantidis, A., Nishino, S., Deisseroth, K., & de Lecea, L. (2010). Tuning arousal with optogenetic modulation of locus coeruleus neurons. Nature Neuroscience, 13(12), 1526–1533. 10.1038/nn.2682

19. Chang, C., Leopold, D. A., Schölvinck, M. L., Mandelkow, H., Picchioni, D., Liu, X., Ye, F. Q., Turchi, J. N., & Duyn, J. H. (2016). Tracking brain arousal fluctuations with fMRI. Proceedings of the National Academy of Sciences, 113(16), 4518–4523. 10.1073/pnas.1520613113

20. Chuang, K.-H., Li, Z., Huang, H. H., Khorasani Gerdekoohi, S., & Athwal, D. (2023). Hemodynamic transient and functional connectivity follow structural connectivity and cell type over the brain hierarchy. Proceedings of the National Academy of Sciences, 120(5), e2202435120. 10.1073/pnas.2202435120

21. Ciric, R., Wolf, D. H., Power, J. D., Roalf, D. R., Baum, G., Ruparel, K., Shinohara, R. T., Elliott, M. A., Eickhoff, S. B., Davatzikos, C., Gur, R. C., Gur, R. E., Bassett, D. S., & Satterthwaite, T. D. (2017). Benchmarking of participant-level confound regression strategies for the control of motion artifact in studies of functional connectivity. NeuroImage, 154, 174–187. 10.1016/j.neuroimage.2017.03.020

22. Fox, M. D., Zhang, D., Snyder, A. Z., & Raichle, M. E. (2009). The Global Signal and Observed Anticorrelated Resting State Brain Networks. Journal of Neurophysiology, 101(6), 3270– 3283. 10.1152/jn.90777.2008

23. Grimm, C., Duss, S. N., Privitera, M., Munn, B. R., Frässle, S., Chernysheva, M., Patriarchi, T., Razansky, D., Wenderoth, N., Shine, J. M., Bohacek, J., & Zerbi, V. (2022). Locus Coeruleus firing patterns selectively modulate brain activity and dynamics (p. 2022.08.29.505672). bioRxiv. 10.1101/2022.08.29.505672

24. Hansen, J. Y., Cauzzo, S., Singh, K., García-Gomar, M. G., Shine, J. M., Bianciardi, M., & Misic, B. (2023). *Integrating brainstem and cortical functional architectures* (p. 2023.10.26.564245). bioRxiv. 10.1101/2023.10.26.564245

25. Hassani, S. A., & Womelsdorf, T. (2023). *Noradrenergic alpha-2a Receptor Stimulation Enhances Prediction Error Signaling in Anterior Cingulate Cortex and Striatum* (p. 2023.10.25.564052). bioRxiv. 10.1101/2023.10.25.564052

26. Hwang, D. Y., Carlezon, W. A., Isacson, O., & Kim, K. S. (2001). A high-efficiency synthetic promoter that drives transgene expression selectively in noradrenergic neurons. Human Gene Therapy, 12(14), 1731–1740. 10.1089/104303401750476230

27. Iannitelli, A. F., & Weinshenker, D. (2023). Riddles in the dark: Decoding the relationship between neuromelanin and neurodegeneration in locus coeruleus neurons. Neuroscience and Biobehavioral Reviews, 152, 105287. 10.1016/j.neubiorev.2023.105287

28. Kelberman, M. A., Rorabaugh, J. M., Anderson, C. R., Marriott, A., DePuy, S. D., Rasmussen, K., McCann, K. E., Weiss, J. M., & Weinshenker, D. (2023). Age-dependent dysregulation of locus coeruleus firing in a transgenic rat model of Alzheimer’s disease. Neurobiology of Aging, 125, 98–108. 10.1016/j.neurobiolaging.2023.01.016

29. Kelberman, M., Keilholz, S., & Weinshenker, D. (2020). What’s That (Blue) Spot on my MRI? Multimodal Neuroimaging of the Locus Coeruleus in Neurodegenerative Disease. Frontiers in Neuroscience, 14, 583421. 10.3389/fnins.2020.583421

30. Lee, J.-Y., You, T., Woo, C.-W., & Kim, S.-G. (2022). Optogenetic fMRI for Brain-Wide Circuit Analysis of Sensory Processing. International Journal of Molecular Sciences, 23(20), Article 20. 10.3390/ijms232012268

31. Li, L., Rana, A. N., Li, E. M., Feng, J., Li, Y., & Bruchas, M. R. (2023). Activity-dependent constraints on catecholamine signaling. Cell Reports, 42(12). 10.1016/j.celrep.2023.113566

32. Liu, T. T., Nalci, A., & Falahpour, M. (2017). The Global Signal in fMRI: Nuisance or Information? NeuroImage, 150, 213–229. 10.1016/j.neuroimage.2017.02.036

33. Liu, X., Zhu, X.-H., Zhang, Y., & Chen, W. (2013). The change of functional connectivity specificity in rats under various anesthesia levels and its neural origin. Brain Topography, 26(3), 363–377. 10.1007/s10548-012-0267-5

34. Liu, Y., Rodenkirch, C., Moskowitz, N., Schriver, B., & Wang, Q. (2017). Dynamic Lateralization of Pupil Dilation Evoked by Locus Coeruleus Activation Results from Sympathetic not Parasympathetic Contributions. Cell Reports, 20(13), 3099–3112. 10.1016/j.celrep.2017.08.094

35. Lu, H., Zou, Q., Gu, H., Raichle, M. E., Stein, E. A., & Yang, Y. (2012). Rat brains also have a default mode network. Proceedings of the National Academy of Sciences, 109(10), 3979–3984. 10.1073/pnas.1200506109

36. Lustberg, D., Iannitelli, A. F., Tillage, R. P., Pruitt, M., Liles, L. C., & Weinshenker, D. (2020). Central norepinephrine transmission is required for stress-induced repetitive behavior in two rodent models of obsessive-compulsive disorder. Psychopharmacology, 237(7), 1973–1987. 10.1007/s00213-020-05512-0

37. Lustberg, D., Tillage, R. P., Bai, Y., Pruitt, M., Liles, L. C., & Weinshenker, D. (2020). Noradrenergic circuits in the forebrain control affective responses to novelty. Psychopharmacology, 237(11), 3337–3355. 10.1007/s00213-020-05615-8

38. Ma, H.-T., Zhang, H.-C., Zuo, Z.-F., & Liu, Y.-X. (2023). Heterogeneous organization of Locus coeruleus: An intrinsic mechanism for functional complexity. Physiology & Behavior, 268, 114231. 10.1016/j.physbeh.2023.114231

39. Majeed, W., Magnuson, M., Hasenkamp, W., Schwarb, H., Schumacher, E. H., Barsalou, L., & Keilholz, S. D. (2011). Spatiotemporal dynamics of low frequency BOLD fluctuations in rats and humans. NeuroImage, 54(2), 1140–1150. 10.1016/j.neuroimage.2010.08.030

40. Marzo, A., Totah, N. K., Neves, R. M., Logothetis, N. K., & Eschenko, O. (2014). Unilateral electrical stimulation of rat locus coeruleus elicits bilateral response of norepinephrine neurons and sustained activation of medial prefrontal cortex. Journal of Neurophysiology, 111(12), 2570–2588. 10.1152/jn.00920.2013

41. Masamoto, K., Fukuda, M., Vazquez, A., & Kim, S.-G. (2009). Dose-dependent Effect of Isoflurane on Neurovascular Coupling in Rat Cerebral Cortex. The European Journal of Neuroscience, 30(2), 242–250. 10.1111/j.1460-9568.2009.06812.x

42. Masamoto, K., & Kanno, I. (2012). Anesthesia and the quantitative evaluation of neurovascular coupling. Journal of Cerebral Blood Flow & Metabolism, 32(7), 1233–1247. 10.1038/jcbfm.2012.50

43. McCall, J. G., Al-Hasani, R., Siuda, E. R., Hong, D. Y., Norris, A. J., Ford, C. P., & Bruchas, M. R. (2015). CRH engagement of the locus coeruleus noradrenergic system mediates stress-induced anxiety. Neuron, 87(3), 605–620. 10.1016/j.neuron.2015.07.002

44. Meyer-Baese, L., Anumba, N., Bolt, T. S., Daley, L., LaGrow, T. J., Zhang, X., Xu, N., Pan, W.-J., Schumacher, E. H., & Keilholz, S. (2024). Variation in the Distribution of Large-scale Spatiotemporal Patterns of Activity Across Brain States (p. 2024.04.26.591295). bioRxiv. 10.1101/2024.04.26.591295

45. Morris, L. S., McCall, J. G., Charney, D. S., & Murrough, J. W. (2020). The role of the locus coeruleus in the generation of pathological anxiety. Brain and Neuroscience Advances, 4, 2398212820930321. 10.1177/2398212820930321

46. Murphy, K., & Fox, M. D. (2017). Towards a consensus regarding global signal regression for resting state functional connectivity MRI. Neuroimage, 154, 169–173. 10.1016/j.neuroimage.2016.11.052

47. Oyarzabal, E. A., Hsu, L.-M., Das, M., Chao, T.-H. H., Zhou, J., Song, S., Zhang, W., Smith, K. G., Sciolino, N. R., Evsyukova, I. Y., Yuan, H., Lee, S.-H., Cui, G., Jensen, P., & Shih, Y.-Y. I. (2021). Chemogenetic activation of Locus Coeruleus Noradrenergic Neurons Modulates the Default Mode Network (p. 2021.10.28.463794). 10.1101/2021.10.28.463794

48. Pagani, M., Gutierrez-Barragan, D., de Guzman, A. E., Xu, T., & Gozzi, A. (2023). Mapping and comparing fMRI connectivity networks across species. Communications Biology, 6(1), Article 1. 10.1038/s42003-023-05629-w

49. Pan, W.-J., Khalizad Sharghi, V., Zhang, X., & Keilholz, S. (2020, August 8). Brain mechanism of anesthesia and sedation: fMRI functional connectivity study with minimized impact of physiological background noise in rats. ISMRM. https://archive.ismrm.org/2020/3958.html

50. Pan, W.-J., Thompson, G. J., Magnuson, M. E., Jaeger, D., & Keilholz, S. (2013). Infraslow LFP correlates to resting-state fMRI BOLD signals. NeuroImage, 74, 288–297. 10.1016/j.neuroimage.2013.02.035

51. Parkes, L., Fulcher, B., Yücel, M., & Fornito, A. (2018). An evaluation of the efficacy, reliability, and sensitivity of motion correction strategies for resting-state functional MRI. NeuroImage, 171, 415–436. 10.1016/j.neuroimage.2017.12.073

52. Pisauro, M. A., Benucci, A., & Carandini, M. (2016). Local and global contributions to hemodynamic activity in mouse cortex. Journal of Neurophysiology, 115(6), 2931–2936. 10.1152/jn.00125.2016

53. Poe, G. R., Foote, S., Eschenko, O., Johansen, J. P., Bouret, S., Aston-Jones, G., Harley, C. W., Manahan-Vaughan, D., Weinshenker, D., Valentino, R., Berridge, C., Chandler, D. J., Waterhouse, B., & Sara, S. J. (2020). Locus coeruleus: A new look at the blue spot. Nature Reviews Neuroscience, 21(11), Article 11. 10.1038/s41583-020-0360-9

54. Privitera, M., Ferrari, K. D., von Ziegler, L. M., Sturman, O., Duss, S. N., Floriou-Servou, A., Germain, P.-L., Vermeiren, Y., Wyss, M. T., De Deyn, P. P., Weber, B., & Bohacek, J. (2020). A complete pupillometry toolbox for real-time monitoring of locus coeruleus activity in rodents. Nature Protocols, 15(8), Article 8. 10.1038/s41596-020-0324-6

55. Prokopiou, P. C., Engels-Domínguez, N., Papp, K. V., Scott, M. R., Schultz, A. P., Schneider, C., Farrell, M. E., Buckley, R. F., Quiroz, Y. T., El Fakhri, G., Rentz, D. M., Sperling, R. A., Johnson, K. A., & Jacobs, H. I. L. (2022). Lower novelty-related locus coeruleus function is associated with Aβ-related cognitive decline in clinically healthy individuals. Nature Communications, 13(1), 1571. 10.1038/s41467-022-28986-2

56. Proudfit, H. K., & Clark, F. M. (1991). The projections of locus coeruleus neurons to the spinal cord. Progress in Brain Research, 88, 123–141. 10.1016/s0079-6123(08)63803-0

57. Raichle, M. E., MacLeod, A. M., Snyder, A. Z., Powers, W. J., Gusnard, D. A., & Shulman, G. L. (2001). A default mode of brain function. Proceedings of the National Academy of Sciences, 98(2), 676–682. 10.1073/pnas.98.2.676

58. Rajkowski, J., Kubiak, P., & Aston-Jones, G. (1994). Locus coeruleus activity in monkey: Phasic and tonic changes are associated with altered vigilance. Brain Research Bulletin, 35(5–6), 607–616. 10.1016/0361-9230(94)90175-9

59. Raut, R. V., Snyder, A. Z., Mitra, A., Yellin, D., Fujii, N., Malach, R., & Raichle, M. E. (2021). Global waves synchronize the brain’s functional systems with fluctuating arousal. Science Advances, 7(30), eabf2709. 10.1126/sciadv.abf2709

60. Saad, Z. S., Gotts, S. J., Murphy, K., Chen, G., Jo, H. J., Martin, A., & Cox, R. W. (2012). Trouble at rest: How correlation patterns and group differences become distorted after global signal regression. Brain Connectivity, 2(1), 25–32. 10.1089/brain.2012.0080

61. Satterthwaite, T. D., Wolf, D. H., Loughead, J., Ruparel, K., Elliott, M. A., Hakonarson, H., Gur, R. C., & Gur, R. E. (2012). Impact of in-scanner head motion on multiple measures of functional connectivity: Relevance for studies of neurodevelopment in youth. NeuroImage, 60(1), 623–632. 10.1016/j.neuroimage.2011.12.063

62. Schwarz, L. A., & Luo, L. (2015). Organization of the Locus Coeruleus-Norepinephrine System. Current Biology, 25(21), R1051–R1056. 10.1016/j.cub.2015.09.039

63. Shine, J. M. (2019). Neuromodulatory Influences on Integration and Segregation in the Brain. Trends in Cognitive Sciences, 23(7), 572–583. 10.1016/j.tics.2019.04.002

64. Smith, S. M., Jenkinson, M., Woolrich, M. W., Beckmann, C. F., Behrens, T. E. J., Johansen-Berg, H., Bannister, P. R., De Luca, M., Drobnjak, I., Flitney, D. E., Niazy, R. K., Saunders, J., Vickers, J., Zhang, Y., De Stefano, N., Brady, J. M., & Matthews, P. M. (2004). Advances in functional and structural MR image analysis and implementation as FSL. NeuroImage, 23, S208–S219. 10.1016/j.neuroimage.2004.07.051

65. Stevens, F. L., Hurley, R. A., Taber, K. H., Hurley, R. A., Hayman, L. A., & Taber, K. H. (2011). Anterior Cingulate Cortex: Unique Role in Cognition and Emotion. The Journal of Neuropsychiatry and Clinical Neurosciences, 23(2), 121–125. 10.1176/jnp.23.2.jnp121

66. Thompson, G. J., Magnuson, M. E., Merritt, M. D., Schwarb, H., Pan, W.-J., McKinley, A., Tripp, L. D., Schumacher, E. H., & Keilholz, S. D. (2013). Short-time windows of correlation between large-scale functional brain networks predict vigilance intraindividually and interindividually. Human Brain Mapping, 34(12), 3280–3298. 10.1002/hbm.22140

67. Tillage, R. P., Foster, S. L., Lustberg, D., Liles, L. C., McCann, K. E., & Weinshenker, D. (2021). Co-released norepinephrine and galanin act on different timescales to promote stress-induced anxiety-like behavior. Neuropsychopharmacology, 46(8), Article 8. 10.1038/s41386-021-01011-8

68. Tillage, R. P., Wilson, G. E., Liles, L. C., Holmes, P. V., & Weinshenker, D. (2020). Chronic Environmental or Genetic Elevation of Galanin in Noradrenergic Neurons Confers Stress Resilience in Mice. Journal of Neuroscience, 40(39), 7464–7474. 10.1523/JNEUROSCI.0973-20.2020

69. Turchi, J., Chang, C., Ye, F. Q., Russ, B. E., Yu, D. K., Cortes, C. R., Monosov, I. E., Duyn, J. H., & Leopold, D. A. (2018). The Basal Forebrain Regulates Global Resting-State fMRI Fluctuations. Neuron, 97(4), 940–952.e4. 10.1016/j.neuron.2018.01.032

70. Uematsu, A., Tan, B. Z., & Johansen, J. P. (2015). Projection specificity in heterogeneous locus coeruleus cell populations: Implications for learning and memory. Learning & Memory, 22(9), 444–451. 10.1101/lm.037283.114

71. Valentino, R. J., & Foote, S. L. (1988). Corticotropin-releasing hormone increases tonic but not sensory-evoked activity of noradrenergic locus coeruleus neurons in unanesthetized rats. The Journal of Neuroscience: The Official Journal of the Society for Neuroscience, 8(3), 1016–1025. 10.1523/JNEUROSCI.08-03-01016.1988

72. van den Berg, M., Adhikari, M. H., Verschuuren, M., Pintelon, I., Vasilkovska, T., Van Audekerke, J., Missault, S., Heymans, L., Ponsaerts, P., De Vos, W. H., Van der Linden, A., Keliris, G. A., & Verhoye, M. (2022). Altered basal forebrain function during whole-brain network activity at pre– and early-plaque stages of Alzheimer’s disease in TgF344-AD rats. Alzheimer’s Research & Therapy, 14(1), 148. 10.1186/s13195-022-01089-2

73. van den Brink, R. L., Pfeffer, T., & Donner, T. H. (2019). Brainstem Modulation of Large-Scale Intrinsic Cortical Activity Correlations. Frontiers in Human Neuroscience, 13. 10.3389/fnhum.2019.00340

74. Vazey, E. M., & Aston-Jones, G. (2014). Designer receptor manipulations reveal a role of the locus coeruleus noradrenergic system in isoflurane general anesthesia. Proceedings of the National Academy of Sciences of the United States of America, 111(10), 3859–3864. 10.1073/pnas.1310025111

75. Waterhouse, B. D., Lin, C. S., Burne, R. A., & Woodward, D. J. (1983). The distribution of neocortical projection neurons in the locus coeruleus. The Journal of Comparative Neurology, 217(4), 418–431. 10.1002/cne.902170406

76. Waterhouse, B. D., Predale, H. K., Plummer, N. W., Jensen, P., & Chandler, D. J. (2022). Probing the structure and function of locus coeruleus projections to CNS motor centers. Frontiers in Neural Circuits, 16, 895481. 10.3389/fncir.2022.895481

77. Watters, H., Fazili, A., Daley, L., Belden, A., LaGrow, T. J., Bolt, T., Loui, P., & Keilholz, S. (2024). Creative tempo: Spatiotemporal dynamics of the default mode network in improvisational musicians (p. 2024.04.07.588391). bioRxiv. 10.1101/2024.04.07.588391

78. Watters, P. A., Martin, F., & Schreter, Z. (1997). Caffeine and Cognitive Performance: The Nonlinear Yerkes–Dodson Law. Human Psychopharmacology: Clinical and Experimental, 12(3), 249–257. 10.1002/(SICI)1099-1077(199705/06)12:3<249::AID-HUP865>3.0.CO;2-J

79. Weinshenker, D. (2018). Long Road to Ruin: Noradrenergic Dysfunction in Neurodegenerative Disease. Trends in Neurosciences, 41(4), 211–223. 10.1016/j.tins.2018.01.010

80. Weinshenker, D., & Holmes, P. V. (2016). Regulation of neurological and neuropsychiatric phenotypes by locus coeruleus-derived galanin. Brain Research, 1641(Pt B), 320–337. 10.1016/j.brainres.2015.11.025

81. Wong, C. W., DeYoung, P. N., & Liu, T. T. (2016). Differences in the resting-state fMRI global signal amplitude between the eyes open and eyes closed states are related to changes in EEG vigilance. NeuroImage, 124(Pt A), 24–31. 10.1016/j.neuroimage.2015.08.053

82. Wong, C. W., Olafsson, V., Tal, O., & Liu, T. T. (2012). Anti-correlated networks, global signal regression, and the effects of caffeine in resting-state functional MRI. NeuroImage, 63(1), 356–364. 10.1016/j.neuroimage.2012.06.035

83. Wong, C. W., Olafsson, V., Tal, O., & Liu, T. T. (2013). The amplitude of the resting-state fMRI global signal is related to EEG vigilance measures. NeuroImage, 83, 983–990. 10.1016/j.neuroimage.2013.07.057

84. Xu, N., Yousefi, B., Anumba, N., LaGrow, T. J., Zhang, X., & Keilholz, S. (2023). QPPLab: A generally applicable software package for detecting, analyzing, and visualizing large-scale quasiperiodic spatiotemporal patterns (QPPs) of brain activity (p. 2023.09.25.559086). bioRxiv. 10.1101/2023.09.25.559086

85. Xu, N., Zhang, L., Larson, S., Li, Z., Anumba, N., Daley, L., Pan, W.-J., Chuang, K.-H., & Keilholz, S. D. (2023). Rodent Whole-Brain fMRI Data Preprocessing Toolbox. Aperture Neuro, 3, 1–3. 10.52294/001c.85075

86. Yan, C.-G., Cheung, B., Kelly, C., Colcombe, S., Craddock, R. C., Di Martino, A., Li, Q., Zuo, X.-N., Castellanos, F. X., & Milham, M. P. (2013). A comprehensive assessment of regional variation in the impact of head micromovements on functional connectomics. NeuroImage, 76, 183–201. 10.1016/j.neuroimage.2013.03.004

87. Yousefi, B., & Keilholz, S. (2021). Propagating patterns of intrinsic activity along macroscale gradients coordinate functional connections across the whole brain. NeuroImage, 231, 117827. 10.1016/j.neuroimage.2021.117827

88. Yousefi, B., Shin, J., Schumacher, E. H., & Keilholz, S. D. (2018). Quasi-Periodic Patterns of Intrinsic Brain Activity in Individuals and their Relationship to Global Signal. NeuroImage, 167, 297–308. 10.1016/j.neuroimage.2017.11.043

89. Yu-Feng, Z., Yong, H., Chao-Zhe, Z., Qing-Jiu, C., Man-Qiu, S., Meng, L., Li-Xia, T., Tian-Zi, J., & Yu-Feng, W. (2007). Altered baseline brain activity in children with ADHD revealed by resting-state functional MRI. Brain and Development, 29(2), 83–91. 10.1016/j.braindev.2006.07.002

90. Zerbi, V., Floriou-Servou, A., Markicevic, M., Vermeiren, Y., Sturman, O., Privitera, M., von Ziegler, L., Ferrari, K. D., Weber, B., De Deyn, P. P., Wenderoth, N., & Bohacek, J. (2019). Rapid Reconfiguration of the Functional Connectome after Chemogenetic Locus Coeruleus Activation. Neuron, 103(4), 702–718.e5. 10.1016/j.neuron.2019.05.034

