## Supplemental Material for "The Effects of Locus Coeruleus Optogenetic Stimulation on Global Spatiotemporal Patterns in Rats"

**SUPPLEMENTARY MATERIAL**

**Supplementary Table 1: Animal Group Sizes by Sex**

| | **6-month WT mCherry** (M/F) | **6-month WT ChR2** (M/F) |
|---|---|---|
| **Baseline** | 4/5 | 2/5 |
| **2 Hz** | 3/5 | 1/3 |
| **5 Hz** | 2/5 | 2/4 |
| **15 Hz Phasic** | 3/5 | 2/4 |

Power Spectral Density Analysis of the Global Signal

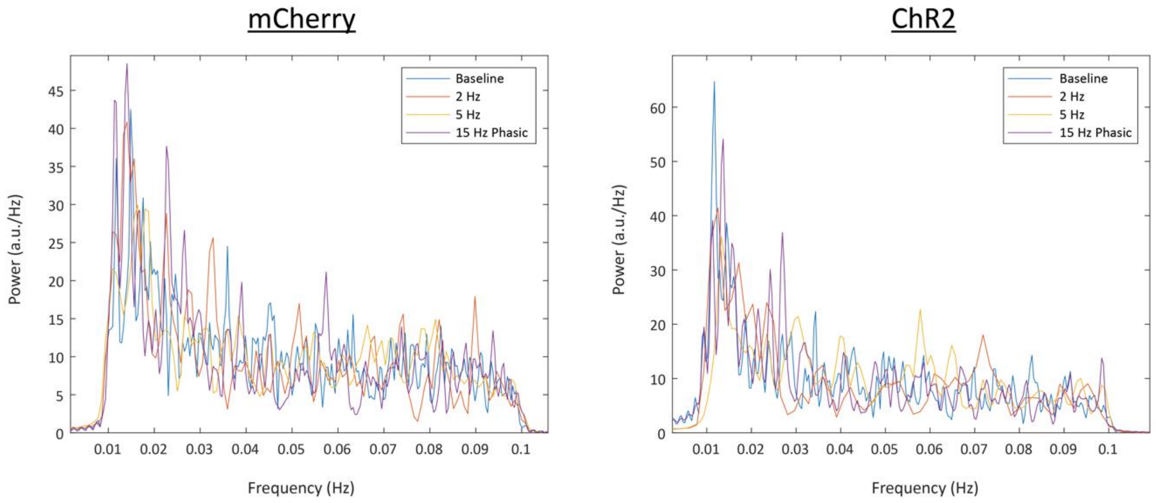


**Figure S1: Power spectral density (PSD) estimates of the global signal.** PSD estimates for the mCherry control animals (left) and the ChR2 stimulated animals (right) are displayed in this figure. All stimulation groups showed a high low-frequency peak in power with a gradual decrease in power as the frequency increased. In the ChR2 animals, the power of this low-frequency peak was higher in the baseline and 15 Hz phasic scans when compared to the control animals. However, no other strong differences were observed between the two groups of animals.

The frequency of the global signal was observed by way of a power spectral density (PSD) analysis. PSD estimates were obtained in MATLAB using Welch's method with a Hamming window and 50% overlap. The global signals for each group were z-scored and concatenated, after which a PSD was calculated for the concatenated signals producing the group results. A PSD estimate was created for the scans at each stimulation level for both the mCherry control and ChR2 stimulated rats (**Figure S1**). Typical for resting-state fMRI scans (Pan et al., 2013), at all stimulation levels a low frequency (< 0.02 Hz) peak in activity was observed, after which the magnitude of power proceeded to steadily taper off at higher frequencies. The magnitude of this low frequency peak was maintained around the same level in the ChR2 stimulated animals when compared to the mCherry controls, with the exception of the baseline and 15 Hz phasic scans for which the low frequency peak was slightly larger in the ChR2 animals. However, no strong notable differences were observed in the power spectra when comparing the corresponding traces for the ChR2 animals to the mCherry controls. In fact, the largest noticeable difference was observed in the baseline scans. It is important to note that this data was bandpass filtered between 0.01 – 1.0 Hz to highlight ultralow fluctuations associated with resting-state activity (Pan et al., 2013). Therefore, it is possible that higher frequencies contained in the global signal may be significantly affected by LC stimulation.

Individual CPCA Incidence

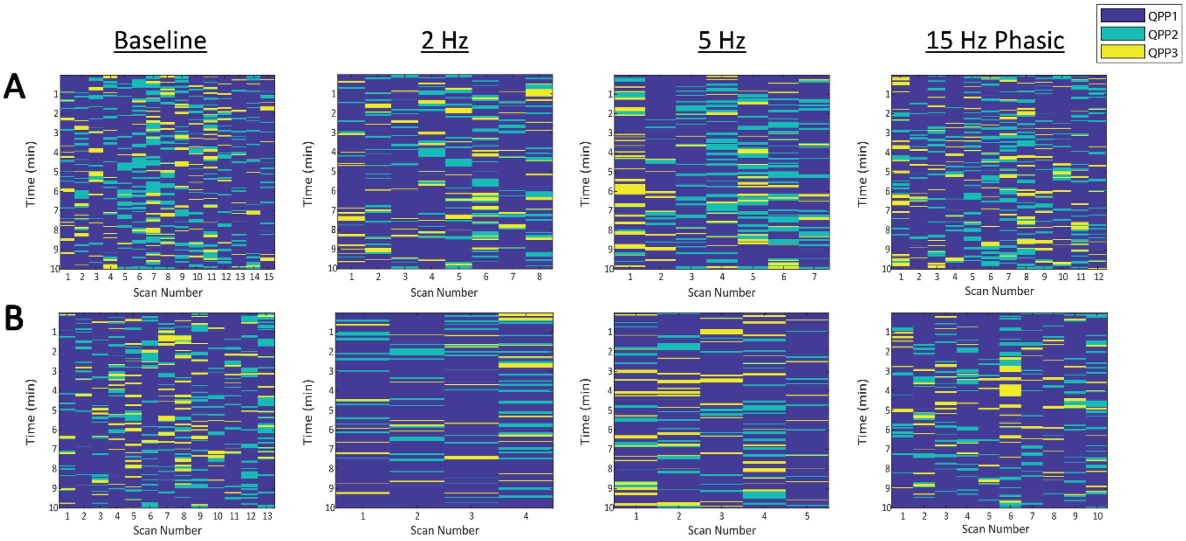


**Figure S2: Incidence of the first three QPPs throughout each scan during LC stimulation.** Each graph shows the number of scans in each group and the principal component that was dominant at each timepoint throughout the scan. The results for the mCherry control animals (A) and ChR2 stimulated animals (B) are shown. These graphs show strong individual variability in terms of timing and the temporal distribution of components across scans and across groups.

The timing of when a component was dominant throughout the scan proved to be highly varied across scans (**Figure S2**). A majority of scan time was dominated by QPP1 with short periods of QPP2 and QPP3 interspersed throughout the scan. Although overall the least amount of time was spent in QPP3, certain scans were seen to spend more time in QPP3 than others (such as scan 1 in the mCherry 5 Hz group and scan 6 in the ChR2 15 Hz phasic group). While longer periods of time (sometimes on the order of minutes) were observed to remain in QPP1, the positioning of these time periods with respect to the beginning and the end of the scan varied quite a bit across rats. No observable pattern as to the timing and sequence of the varying component states was seen based on virus or stimulation level.
